# Affinity-optimizing variants within cardiac enhancers disrupt heart development and contribute to cardiac traits

**DOI:** 10.1101/2022.05.27.493636

**Authors:** Granton A Jindal, Alexis T Bantle, Joe J Solvason, Jessica L Grudzien, Agnieszka D’Antonio-Chronowska, Fabian Lim, Sophia H Le, Reid O Larsen, Adam Klie, Kelly A Frazer, Emma K Farley

## Abstract

Enhancers direct precise gene expression patterns during development and harbor the majority of variants associated with disease. We find that suboptimal affinity ETS transcription factor binding sites are prevalent within *Ciona* and human developmental heart enhancers. Here we demonstrate in two diverse systems, *Ciona intestinalis* and human iPSC-derived cardiomyocytes (iPSC-CMs), that single nucleotide changes can optimize the affinity of ETS binding sites, leading to gain-of-function gene expression associated with heart phenotypes. In *Ciona*, ETS affinity-optimizing SNVs lead to ectopic expression and phenotypic changes including two beating hearts. In human iPSC-CMs, an affinity-optimizing SNV associated with QRS duration occurs within an SCN5A enhancer and leads to increased enhancer activity. Our mechanistic approach provides a much-needed systematic framework that works across different enhancers, cell types and species to pinpoint causal enhancer variants contributing to enhanceropathies, phenotypic diversity and evolutionary changes.

**In Brief:** The prevalent use of low-affinity ETS sites within developmental heart enhancers creates vulnerability within genomes whereby single nucleotide changes can dramatically increase binding affinity, causing gain-of-function enhancer activity that impacts heart development.

**Highlights:**

ETS affinity-optimizing SNVs can lead to migration defects and a multi-chambered heart.
An ETS affinity-optimizing human SNV within an SCN5A enhancer increases expression and is associated with QRS duration.
Searching for ETS affinity-optimizing variants is a systematic and generalizable approach to pinpoint causal enhancer variants.

## Introduction

Enhancers are genomic elements that act as switches to control the location, level, and timing of gene expression to ensure the successful development and integrity of an organism (Levine, 2010). Sequence changes within enhancers can lead to changes in gene expression, phenotypic diversity, evolutionary changes, and disease (Lettice et al., 2008; Smemo et al., 2014; Tournamille et al., 1995). Indeed, most disease-associated variants within the genome are thought to lie within enhancers (Maurano et al., 2012; Tak and Farnham, 2015; Visel et al., 2009). Yet, pinpointing causal base-pair changes within enhancers is exceptionally challenging as they are typically embedded within a sea of inert variants (Gallagher and Chen-Plotkin, 2018). Our inability to systematically predict causal enhancer variants is stalling efforts to harness the full potential of genomic data to understand the genetic basis of disease and diagnose and treat patients. Here we use a systematic and generalizable approach to pinpoint causal enhancer variants within the context of heart development across chordates.

We and others have previously shown that developmental enhancers contain suboptimal affinity sites (also referred to as submaximal and low-affinity) which are important for tissue-specific gene expression (Crocker et al., 2015; Delker et al., 2019; Farley et al., 2015, 2016; Kribelbauer et al., 2019; Swanson et al., 2011). Suboptimal affinity binding sites for transcriptional activators ensure that the enhancer is only active where all factors are present at the right concentration, thus ensuring combinatorial control of gene expression. Increasing the affinities of all sites within an enhancer results in loss of combinatorial control: the enhancer no longer exhibits tissue-specific expression but rather is active in all cells that receive even low levels of the transcriptional activator (Crocker et al., 2015; Farley et al., 2015). In addition to ensuring tissue specificity, suboptimal affinity binding sites are also likely to be important for ensuring the correct levels of expression within a particular cell type. While the use of suboptimal affinity binding sites to encode precise gene expression has been seen in a variety of contexts, it has not yet been explored within heart development across chordates.

Given the prevalence of low-affinity binding sites within developmental enhancers, we hypothesize that single nucleotide variants (SNVs) that increase the affinity of binding sites could cause gain-of-function gene expression that contributes to phenotypic change. Variants that alter binding affinity have been linked to changes in gene expression and phenotypes in a handful of cases, however these are one off examples identified in an ad hoc way (Bond et al., 2004; French et al., 2013; Grant et al., 1996; Huang et al., 2014; Jia et al., 2009; Tuupanen et al., 2009). In contrast, here we investigate if searching for affinity-optimizing variants is a generalizable strategy to pinpoint enhancer variants that contribute to gain-of-function gene expression and phenotypic changes within the context of heart development.

Heart development is a highly conserved process in which FGF (Fibroblast Growth Factor) signaling is critical for specification and migration of heart cells across bilaterians (Harvey, 2002; Zaffran and Frasch, 2002). FGF signaling mediates changes in gene expression through binding of activated ETS transcription factors to enhancers (Sharrocks, 2001). Loss and gain of FGF, ETS, and other members of this signaling pathway cause heart defects in organisms as diverse as *Drosophila* and human (Beiman et al., 1996; Buckingham et al., 2005; Reifers et al., 2000). For example, gain of FGF is implicated in cardiac hypertrophy (House et al., 2010) and manipulation of ETS-1 in mice recapitulates some of the most common congenital heart defects in Jacobson syndrome (Ye et al., 2010). Gain of ETS expression is also associated with evolution of a multichambered heart (Davidson et al., 2006). Here we focus on systematically pinpointing SNVs within enhancers regulated by FGF signaling that could alter gene expression, cell behavior, and heart function.

*Ciona intestinalis type A*, also known as *Ciona robusta*, (*Ciona*) is a marine chordate and member of the urochordates, the sister group to vertebrates (Delsuc et al., 2006). Like the vertebrate heart, *Ciona* has a pumping heart within a pericardium. Additionally in both vertebrates and *Ciona* specification of the heart involves ETS and MesP (Davidson, 2007). Migration of the heart cells to the ventral midline is dependent on FGF signaling and occurs upon activation of the gene FoxF in the Trunk Ventral Cells (TVCs) which give rise to the heart and pharyngeal cells (Beh et al., 2007; Christiaen et al., 2008). For simplicity, we will refer to the TVCs as the heart precursor cells. *Ciona* is experimentally tractable allowing us to investigate enhancer activity in hundreds of developing embryos (Davidson, 2007). Furthermore, the migration of the heart precursor cells to the ventral midline is easily visualized allowing us to study the impact of changes in the FoxF enhancer and FoxF expression on migration and heart development (Davidson, 2007). Thus, *Ciona* provides an ideal system to investigate the relationship between enhancer sequence, tissue-specific gene expression and phenotypes.

Here we find, like other developmental enhancers (Crocker et al., 2015; Farley et al., 2015, 2016), the FoxF enhancer contains incredibly low-affinity ETS binding sites that are necessary to restrict expression of FoxF to the heart precursor cells (Figure 1A). A major consequence of low-affinity sites within functional enhancers is that single base-pair changes that violate the regulatory principle of suboptimization can dramatically increase the affinity of ETS binding sites. In the most extreme case, we observe that a binding site can go from 0.12 binding to 0.97 binding relative to consensus with a single nucleotide change. The higher affinity ETS sites cause ectopic, gain-of-function enhancer activity in non-heart cells that contain lower levels of activated ETS (Figure 2). This can lead to abnormal migration of these non-heart cells to the ventral midline (Figure 3) and defects in heart development, such as enlarged hearts or even two beating hearts (Figure 4).

**Figure 1.**
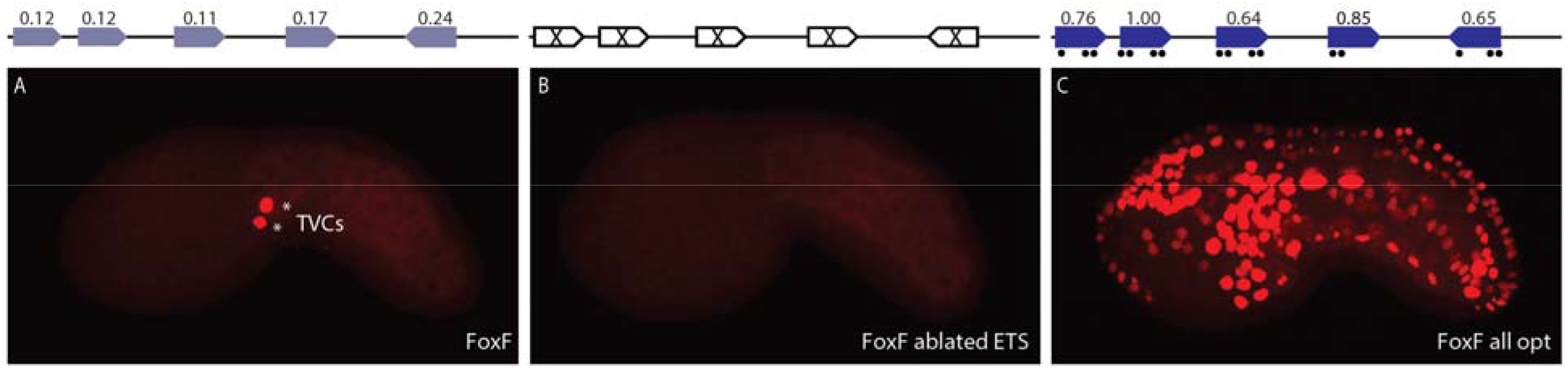
Very low-affinity ETS sites are necessary for tissue-specific expression. **A.** The WT FoxF enhancer drives expression in the heart precursor cells (TVCs). **B.** An enhancer with all five ETS sites ablated (FoxF ablated ETS) drives no enhancer activity. **C.** Optimizing the affinity of all ETS sites (FoxF all opt) leads to ectopic expression in all cell types receiving FGF signaling including: TVCs, ATMs, endoderm, mesenchyme, notochord, and nervous system. The relative affinity can be seen above each site; below each site, dots mark the nucleotide changes relative to WT. Note that ETS6 was also ablated in FoxF -all ETS. ETS6 is not shown in the schematic as ablation of this site did not impact expression (see Figure S2 for more information). Heart precursor cells (TVCs) are marked by asterisks in panel A.

**Figure 2.**
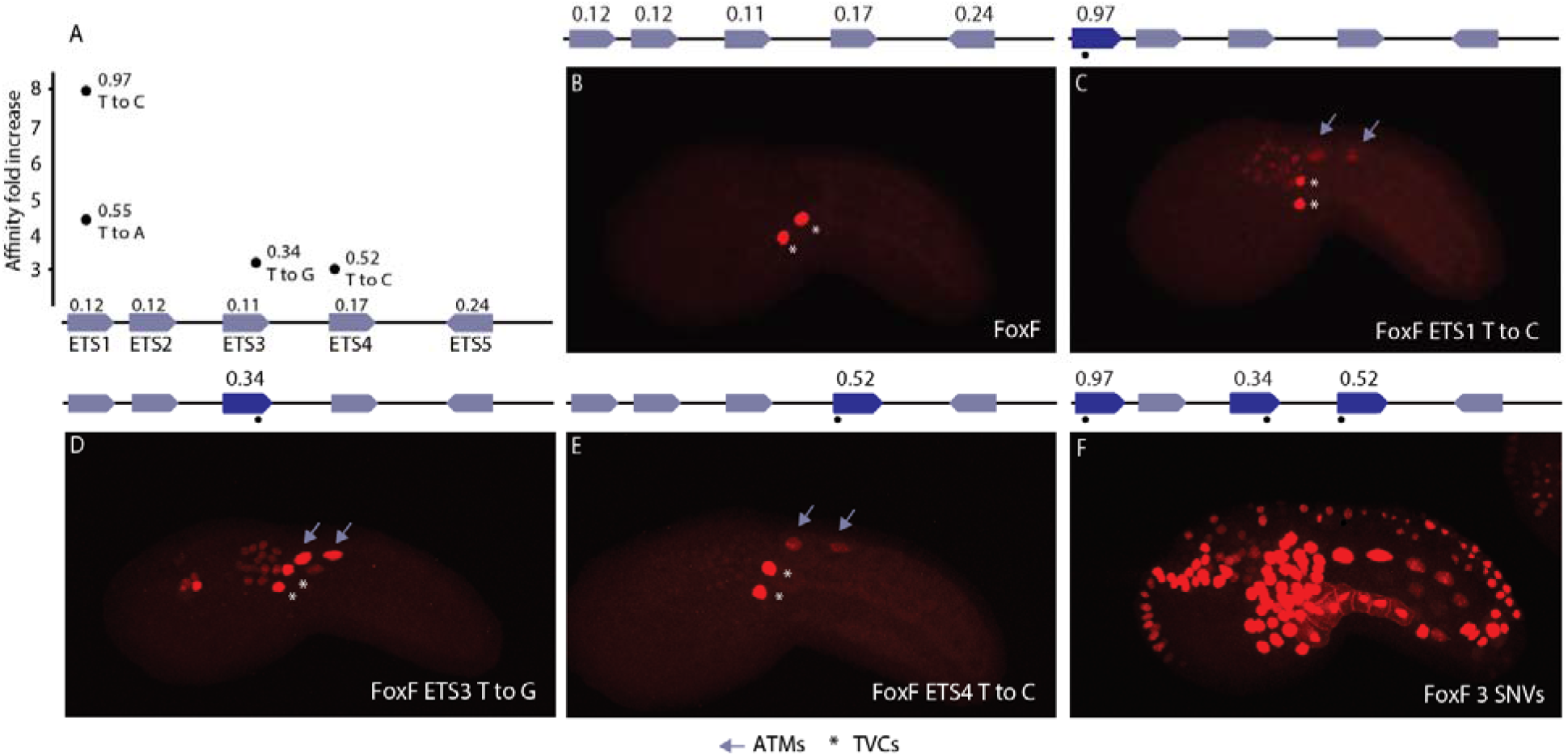
Affinity-optimizing SNVs lead to loss of tissue-specific expression. **A.** Schematic of FoxF enhancer showing affinity-optimizing SNVs. Dots show affinity-optimizing SNVs and fold increase. Starting affinity and final affinities are shown. **B.** The WT FoxF enhancer drives expression in the heart precursor cells (TVCs). **C.** The FoxF ETS1 T to C enhancer harbors a single nucleotide change in the ETS1 site which creates an 8-fold increase in affinity. This enhancer drives expression in the TVCs, ATMs, and mesenchyme. **D.** The FoxF ETS3 T to G enhancer, in which a single base-pair change in ETS3 leads to a 3-fold increase in affinity, drives expression in the TVCs, ATMs, mesenchyme, and neural tissues. **E.** The FoxF ETS4 T to C enhancer, where a single base-pair change in the ETS4 site leads to a 3-fold increase in affinity, drives expression in the TVCs and ATMs. **F.** The FoxF 3 SNVs enhancer contains three affinity-optimizing SNVs and drives expression in many cell types receiving FGF signaling. Heart precursor cells (TVCs) are marked by asterisks in all panels except F. Anterior Tail Muscle cells (ATMs) are marked with arrows in all panels except F. Above each image is a schematic of the enhancer with ETS binding sites shown, values are affinities, dots mark SNVs.

**Figure 3.**
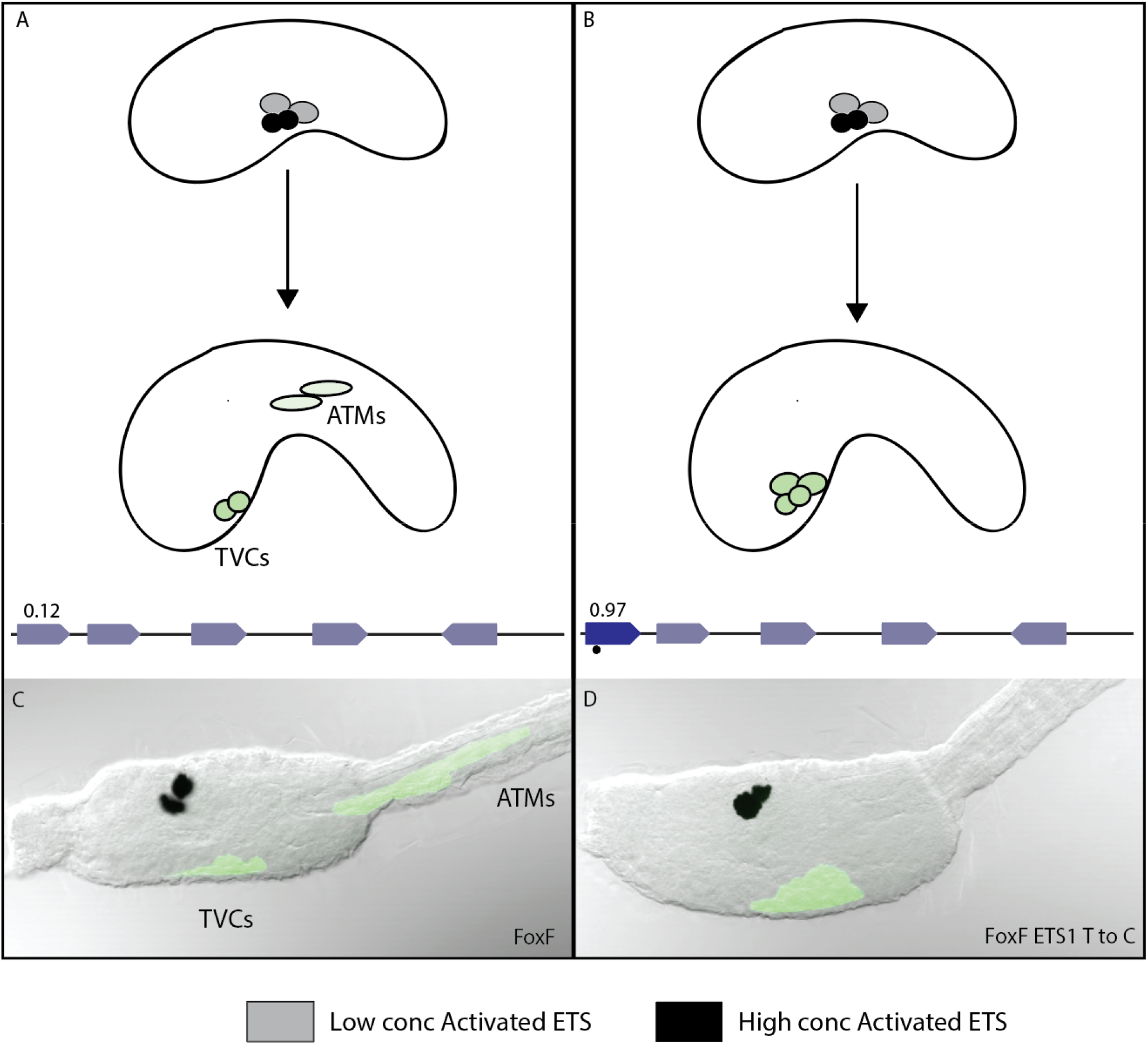
Affinity-optimizing SNVs cause migration defects. **A.** Schematic showing a tailbud wildtype embryo. **B.** Schematic showing a tailbud embryo containing the FoxF ETS1 T to C enhancer driving FoxF. TVC cells have high levels of activated ETS, while ATMs have low levels of activated ETS. In later stage embryos, the FoxF enhancer drives expression of FoxF in the TVCs; this causes them to migrate to the ventral midline **(A)**. In later stage embryos with the FoxF ETS1 T to C enhancer driving FoxF, the ATMs express FoxF and migrate with the TVCs to the ventral midline **(B)**. **C.** Embryo with WT FoxF enhancer driving FoxF, co-electroporated with a MesP enhancer driving GFP that marks the ATMs and TVCs at 16 hpf; GFP expression can be seen in the TVCs and the ATMs. **D.** Embryo with FoxF ETS1 T to C enhancer driving FoxF, co-electroporated with a MesP enhancer driving GFP at 16 hpf; all cells migrate to the ventral midline. Black cells in **C** and **D** are the otolith and ocellus, pigmented cells in the anterior sensory vesicle.

**Figure 4.**
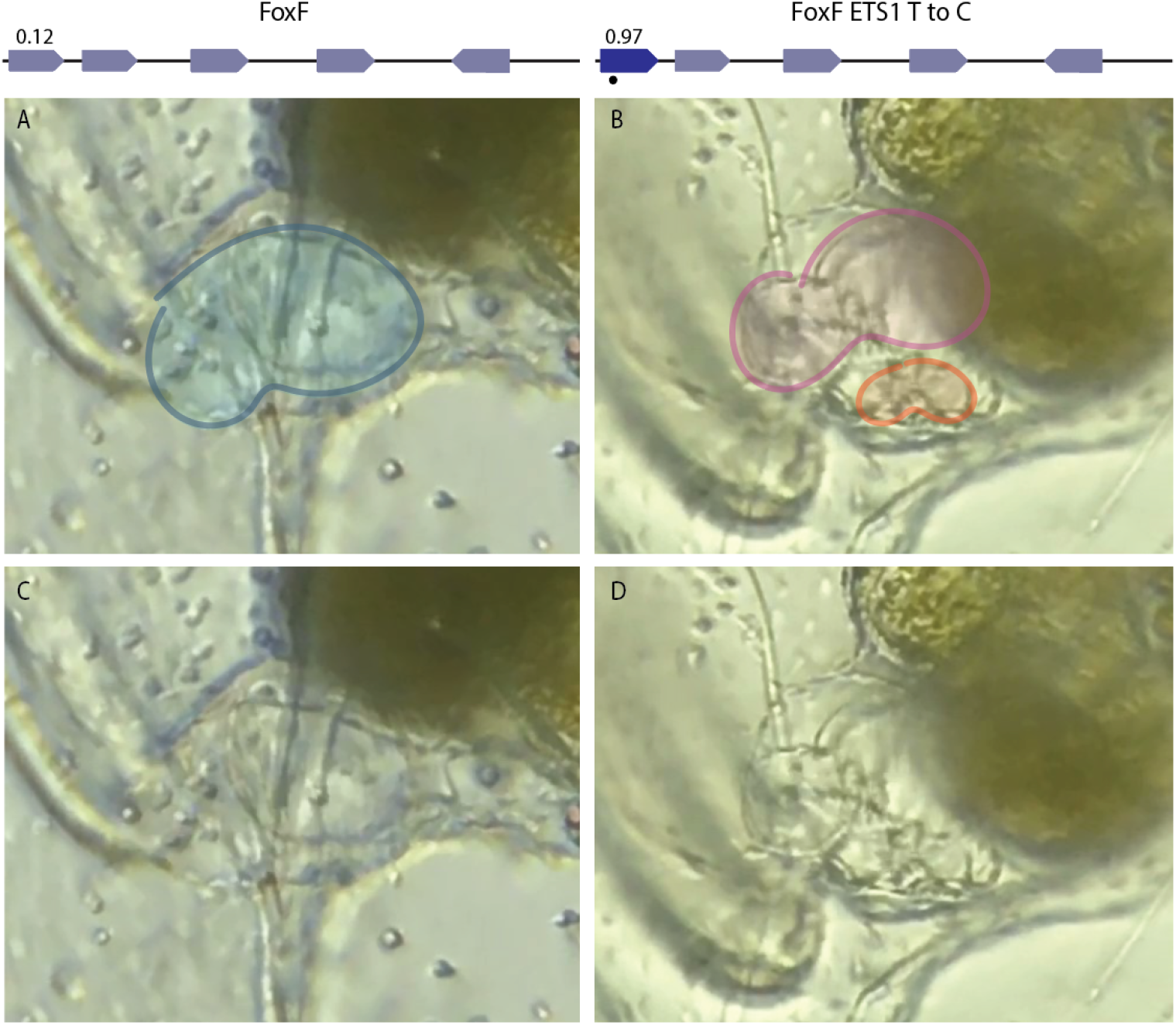
Affinity-optimizing single base-pair mutations disrupt heart development. **A.** Juvenile with WT FoxF enhancer driving FoxF with heart colored in blue; normal heart morphology is seen. **B.** Juvenile with FoxF ETS1 T to C enhancer driving FoxF with main heart colored in pink, auxiliary heart colored in orange; abnormal heart morphology is seen in 79% (41/52) of embryos with migration defects. Hearts were false colored digitally. **C** and **D** are original, uncolored versions of **A** and **B** respectively. Schematic of enhancers with base-pair change marked with dot shown at top.

The FoxF enhancer is not an anomaly. Indeed, we find that the majority of putative enhancers within *Ciona* and human developing heart have low-affinity ETS sites and that affinity-optimizing SNVs occur within these enhancers (Figure 5). We pinpoint 446 ETS affinity-optimizing SNVs that are associated with cardiac traits. 15% of these are within mapped developmental human enhancers making them excellent candidates for causality. One cardiac trait that we focus on is QRS duration. The duration of the QRS complex on a resting, standard 12-lead electrocardiogram (ECG) represents the electrical depolarization of the ventricles as an impulse travels through the cardiac conduction system and the ventricular myocardium. Delay in cardiac ventricular conduction results in increased QRS duration and predicts heart failure prognosis, sudden death, and cardiovascular (CV) mortality in patients with and without left ventricular dysfunction, independent of traditional CV risk factors (Swenson et al., 2019).

**Figure 5.**
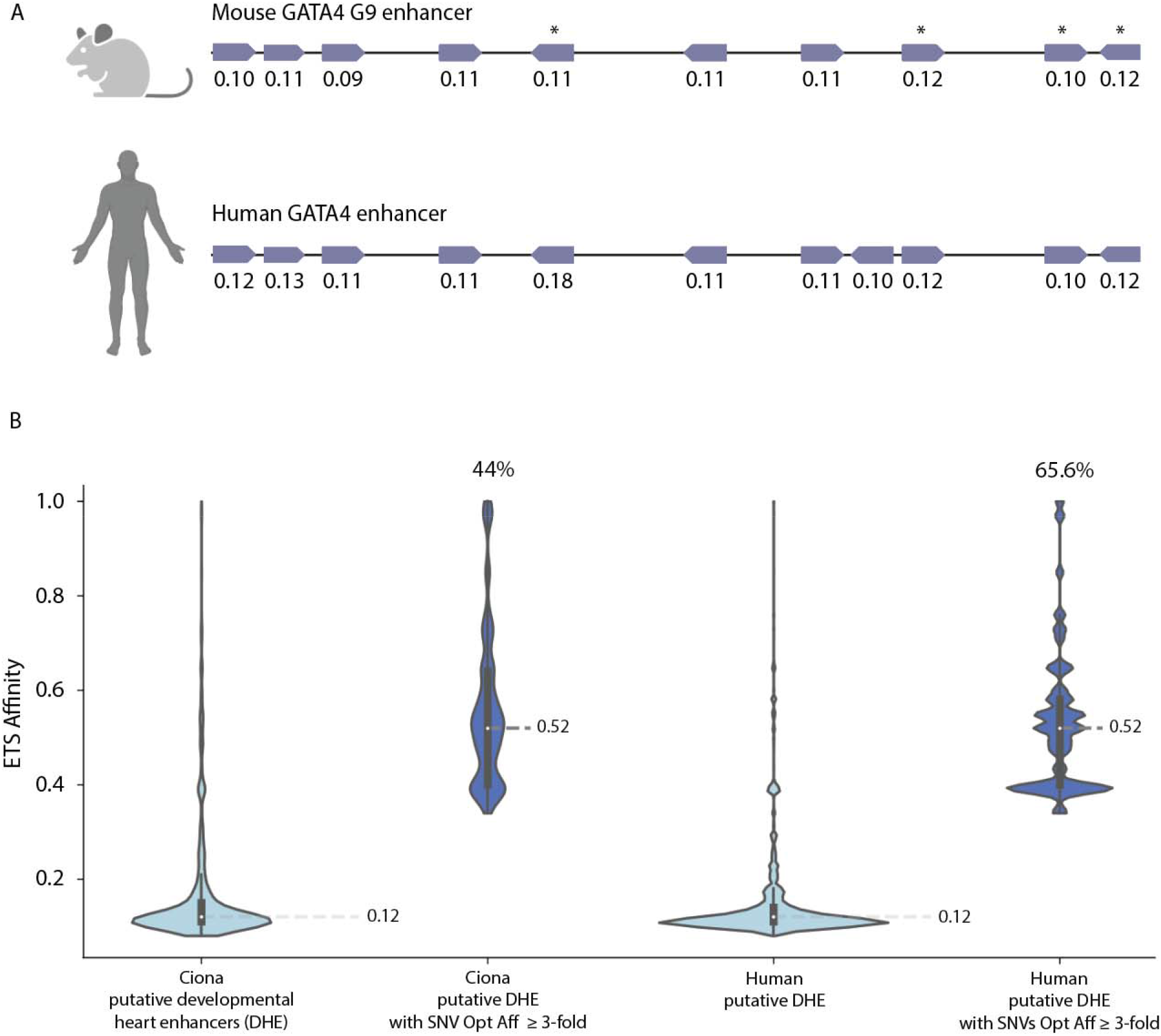
Suboptimal affinity ETS binding sites are prevalent within *Ciona*, mouse and human enhancers. The majority of these ETS sites can be optimized by a SNV. **A.** Schematics of human and mouse GATA4 enhancers showing high conservation of low-affinity ETS sites. Asterisks mark functionally validated ETS sites in mouse (Schachterle et al., 2012). **B.** Violin plot of ETS sites within *Ciona* (left) and human (right) putative developmental heart enhancers (DHE) identified from ATAC-seq (*Ciona*) and ATAC-seq and ChIP-seq for p300 and H3K27Ac (human). 44% (6,615) of *Ciona* DHE contain an ETS site optimizable ≥ 3-fold. 65.6% (129,882/197,963) of human DHE contain an ETS site optimizable ≥ 3-fold.

One of the most highly associated QRS duration GWAS loci is the SCN5A locus (Sotoodehnia et al., 2010). SCN5A is an ion channel that is critical for correct conduction of the heart (Veerman et al., 2015). Both gain- and loss-of-function variants within the coding region of SCN5A, and a noncoding variant, lead to changes in SCN5A function and subsequently heart conduction (Li et al., 2018; Veerman et al., 2015). Overexpression of SCN5A in mice leads to changes in heart conduction (Liu et al., 2015; Zhang et al., 2007). We identify a variant associated with QRS duration within an SCN5A enhancer that leads to an 8-fold increase in ETS binding affinity and a significant increase in expression when tested in human iPSC-derived cardiomyocytes (iPSC-CMs) (Figure 6). This indicates that this variant may lead to gain-of-function expression of SCN5A and alterations in heart conduction. Collectively, our studies across *Ciona* and human developmental heart enhancers highlight that the prevalent use of low-affinity binding sites to encode precise gene expression creates a vulnerability within genomes whereby affinity-optimizing variants can lead to gain-of-function gene expression that contribute to cardiac phenotypes.

**Figure 6.**
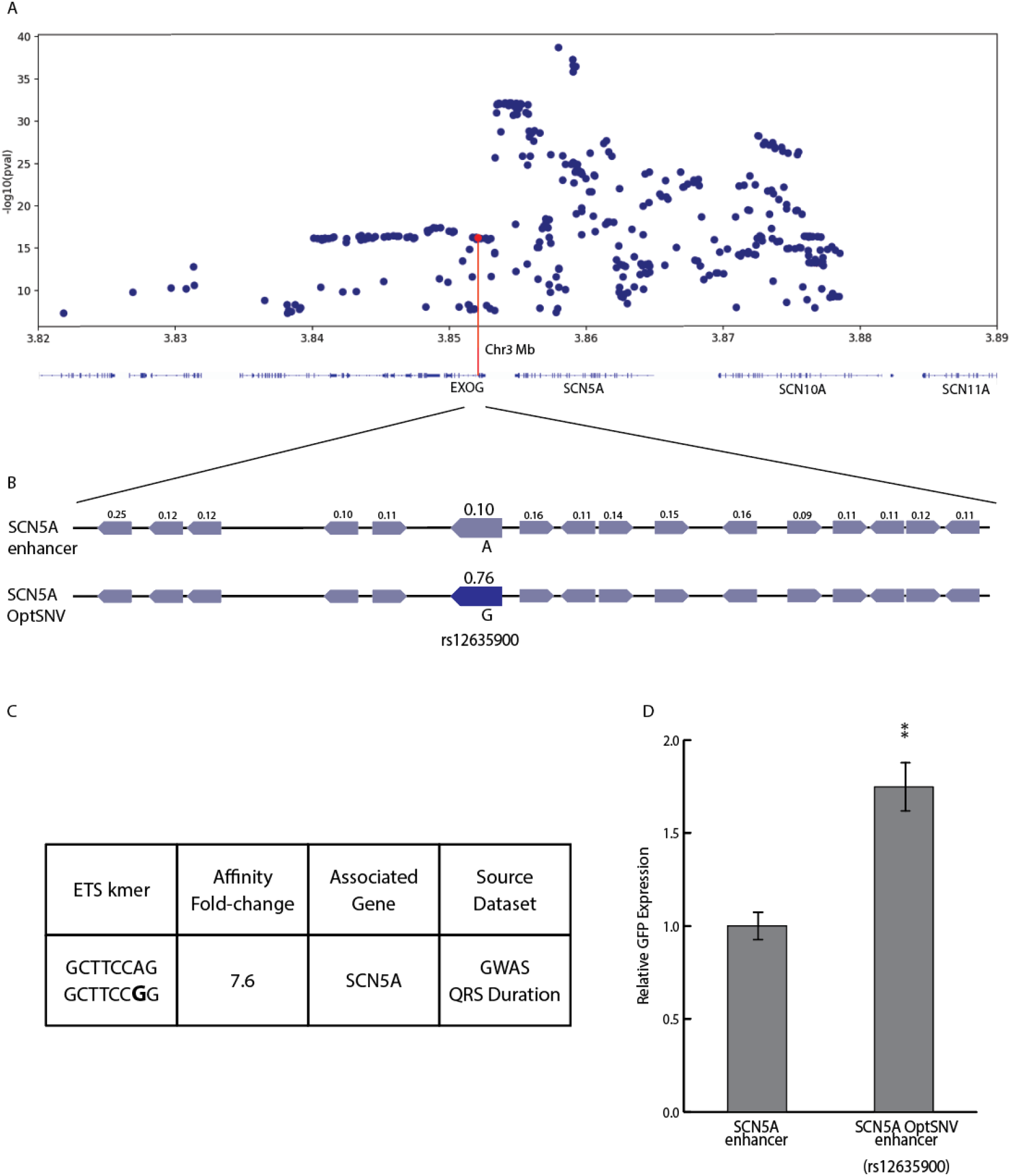
Affinity-optimizing SNV associated with QRS duration occurs within an SCN5A enhancer and drives gain-of-function gene expression within human iPSC-derived cardiomyocytes. **A.** Plot showing all statistically significant SNVs associated with QRS duration within the SCN5A locus with logarithmically scaled P-value on the y-axis and genomic position on chromosome 3 on the x-axis. The ETS affinity-optimizing SNV is shown in red. **B.** Schematics represent enhancer region used for reporter assays within the fetal human heart H3K27Ac peak. There are 16 low-affinity ETS sites annotated. The ETS-optimizing SNV is also shown. **C.** Table with details of SCN5A SNV including ETS 8-mer, affinity change, associated gene, and source dataset. **D.** qPCR data from reporter assay within iPSC-derived human cardiomyocytes. The ETS-optimizing SNV results in a 1.8-fold increase in GFP expression relative to reference SCN5A enhancer. Error bars represent standard deviation of two replicates. ** = P-value < 0.005 (p-value determined by Two-tailed Student’s T-test).

## Results

### The FoxF enhancer drives expression in heart precursor cells and contains suboptimal affinity ETS binding sites

The FoxF enhancer drives expression in the trunk ventral cells (TVCs), which give rise to the heart and pharyngeal cells (Wang et al., 2019). For simplicity, we will refer to these cells as heart precursor cells. The FoxF enhancer activates expression in the heart precursor cells in response to activated ETS (Fig 1A). Three functional ETS sites within this enhancer have been identified (Beh et al., 2007). To identify additional ETS sites within the FoxF enhancer, we searched for ETS cores (GGAW) within the enhancer. We identified three additional putative ETS sites; affinities of all six ETS sites were measured using Protein Binding Microarray data (PBM), which measures the interaction of ETS with all possible 8-mers (Berger and Bulyk, 2009; Wei et al., 2010). As we have done in previous publications, we use PBM data for mouse ETS-1 as a proxy for *Ciona* ETS-1 binding as the DNA binding domain and binding specificity of ETS-1 is highly conserved in species as diverse as *Drosophila, Ciona*, mouse and human (Farley et al., 2015; Nitta et al., 2015) (Figure S1). Using PBM datasets, the highest affinity binding 8-mers has a score of 1.0 or 100%. In *Drosophila, Ciona*, mouse, and human, the highest affinity 8mer is CCGGAAGT (Farley et al., 2015; Nitta et al., 2015; Wei et al., 2010). The relative affinity is calculated by comparing binding measurements between the highest affinity sites with all other 8-mers. Strikingly, we find that the putative ETS sites found within the FoxF enhancer all bind ETS with incredibly low relative affinities of 0.09 to 0.24 (Figures 1A, S2).

### Suboptimal affinity binding sites are necessary for tissue-specific expression

We first sought to determine if these six low-affinity ETS sites were necessary for expression within the heart precursor cells. ETS sites were ablated by introduction of a single base-pair change that disrupts hydrogen bonding between the ETS transcription factor and the DNA motif (Schachterle et al., 2012). We also ensured that these ablations did not create ATTA sites or Ebox motifs. Ablating all of the ETS sites in parallel (FoxF -all ETS) leads to complete loss of enhancer activity (Figures 1B, S2). Having seen the impact of ablating all sites, we next wanted to investigate the individual contribution of each ETS site to FoxF enhancer activity. To do this, we ablated individual ETS sites by point mutations. Ablating individual sites lead to a significant reduction in expression for five of the six ETS sites (Figure S2). The only site without a reduction in expression upon ablation was ETS6. While loss of this site doesn’t lead to reduction of expression, we cannot rule out that this site is redundantly involved in enhancer activity (Fuqua et al., 2020; Lettice et al., 2012; Spivakov, 2014). Together, these results demonstrate that five incredibly suboptimal affinity ETS sites are necessary for the expression of the FoxF enhancer in the heart precursor cells.

### Enhancer grammar likely influences the role of each ETS site

Interestingly, ablating the ETS5 site, which has the highest affinity (0.24), did not have the strongest impact on expression (Figure S2). Instead, ablation of the lowest affinity functional site, ETS3 (0.11 affinity), has the largest impact on enhancer activity (Figure S2). Indeed, ETS3 is the ETS site closest to an ATTA motif and an Ebox binding site, which are both known to be important for FoxF expression (Figure S3) (Beh et al., 2007; Woznica et al., 2012). The significant reduction in enhancer expression upon ablation of this low-affinity ETS site suggests that the combinatorial interaction of this low-affinity ETS site with the ATTA and Ebox sites likely influences the role of this ETS3 sites and its contribution to gene expression. These results suggest that the higher affinity sites are not necessarily the most functionally significant and hint at the importance of enhancer grammar (Jindal and Farley, 2021).

### Suboptimal affinity ETS sites are required for tissue-specific expression

To see if the low-affinity ETS sites within the FoxF enhancer are required for tissue-specific expression, we increased the affinity of the five ETS sites necessary for activity. We altered these such that we kept the core GGAA or GGAT constant and did not create de novo ETS sites, Ebox or ATTA sites. Adhering to these constraints, we were able to optimize the affinity of the five necessary sites to 0.76, 1.0, 0.64, 0.85, and 0.65, respectively. Optimizing the affinity of all five ETS sites (FoxF all ETS opt) within the enhancer leads to extremely ectopic expression in all tissues receiving FGF signaling, including the anterior tail muscle cells (ATMs), mesenchyme, and neural tissues (Figures 1C, S4). Thus, the use of extremely low-affinity sites (with a median affinity of 0.12) is important for maintaining heart-specific expression of the FoxF enhancer. While low-affinity sites have been identified in other enhancers, the sites within the FoxF enhancer are lower affinity than previously documented ETS sites in the neural tissues and notochord enhancers (Farley et al., 2015, 2016). Increasing the affinity of these sites converts a tissue-specific enhancer into an enhancer active in every cell type where FGF signaling occurs. Similar results have been seen upon optimizing the affinity of sites within neural and notochord enhancers (Farley et al., 2015, 2016).

### Single nucleotide variants can dramatically increase the ETS binding affinity

Enhancers are thought to harbor the majority of mutations associated with disease and many of these are single nucleotide variants (SNVs) (Maurano et al., 2012; Tak and Farnham, 2015; Visel et al., 2009). Having seen that optimizing the affinity of all ETS sites leads to loss of tissue-specific expression, we wondered if any single base-pair changes could optimize the affinity of ETS binding sites. We made every possible single base-pair change within the ETS sites of the FoxF enhancer *in silico* and measured the resulting affinity change. We were struck by how easy it was to increase the affinity of an ETS site with a single base-pair change. Three of the five ETS sites can have their affinity increased by ≥3-fold by a single base-pair change. Two of these affinity-optimizing variants occur in the ETS1 site, while the other two affinity-optimizing SNVs occur in ETS3 and ETS4 sites (Figure 2A).

### Affinity-optimizing SNVs lead to gain-of-function enhancer activity in non-heart cells

The greatest affinity change occurs within the ETS1 site, where a single base-pair change increases the affinity 8-fold, creating an almost consensus site, with an affinity of 0.97. When we make this nucleotide change within the FoxF enhancer (FoxF ETS1 T to C), the enhancer drives ectopic expression in the ATMs and mesenchyme, both cell types of which receive FGF signaling (Figures 2A, C). Notably, this single base-pair change does not significantly change expression within the heart precursor cells (TVCs) but leads to expression in the ATMs in over 80% of embryos (Figures 2C, S4).

Even more subtle, yet still dramatic, increases in affinity are seen within the ETS3 and ETS4 sites. A single base-pair change within the ETS3 site causes a 3-fold increase in binding affinity and changes the affinity from 0.11 to 0.34. This affinity-optimizing SNV causes >60% of embryos to have ATM expression and 40% of embryos to have neural expression (Figures S4, 2D). The affinity-optimizing SNV within the ETS4 site changes the affinity from 0.17 to 0.52. This 3-fold increase in ETS affinity causes ectopic expression in the ATMs, mesenchyme, and neural cells (Figures 2E, S4). These results suggest that affinity-optimizing SNVs, and even ones that upon optimization are still relatively low-affinity sites, can lead to loss of tissue-specific gene expression.

It is thought that the majority of enhancer variants that contribute to changes in gene expression are thought to have small effect sizes and that disruption of development and cellular identity occurs due to the combination of many variants (Boyle et al., 2017; Fisher, 1918; Frazer et al., 2009; Mathieson, 2021). We therefore wondered what impact all three affinity-optimizing SNVs in combination would have on gene expression. The FoxF 3 SNVs enhancer drives expression in many tissues responding to FGF signaling and thus the effect of these three SNVs appears to be synergistic (Figure 2F and S4).

### Affinity-optimizing SNVs lead to migration defects

We next wished to understand how these affinity-optimizing variants would impact cellular behavior. All three enhancers containing an optimizing SNV show ectopic expression in the ATMs (Figure 2). ATMs and TVCs are related cell types; however, the TVCs contain higher levels of activated ETS due to an enrichment of FGF receptors on these cells (Cota and Davidson, 2015). The high levels of activated ETS within the TVCs causes TVC-specific expression of FoxF and migration of the heart precursor cells (TVCs) to the ventral midline (Cooley et al., 2011). As the ATMs have low levels of activated ETS, they do not express FoxF and do not migrate (Figure 3 and Movie S1) (Cooley et al., 2011). Previous studies have shown that ectopic expression of FoxF protein or overexpression of constitutively active ETS in the ATMs causes the ATMs to migrate with the TVCs to the ventral midline (Beh et al., 2007; Davidson et al., 2006). We wondered if the FoxF ETS1 T to C base-pair change that leads to an 8-fold increase in ETS affinity and ectopic expression in the ATMs would be sufficient to cause migration of the ATMs to the ventral midline.

The majority of variants contributing to phenotypic diversity and disease lie in the non-coding genome (Maurano et al., 2012; Tak and Farnham, 2015; Visel et al., 2009). However, in most cases each non-coding variant is thought to contribute a small effect (Boyle et al., 2017; Fisher, 1918; Frazer et al., 2009). It is the combination of many variants of small effect size that lead to phenotypic changes. This makes identifying enhancer variants that contribute to phenotypes challenging. In order to see if affinity-optimizing SNVs have the potential to contribute to phenotypic changes, we used the 3kb regulatory element for FoxF including the endogenous promoter to drive FoxF expression in developing embryos. When the FoxF gene is driven by the wild-type FoxF enhancer sequence, we observe no migration of ATMs to the ventral midline (Figures 3C, S5 and Movie S1) and normal hearts are observed at the juvenile stage (Movie S2). However, when we drive expression of the FoxF gene with the FoxF ETS1 T to C enhancer, we see migration of the ATMs into the trunk in 9.3% of embryos (Figures 3, S5, Movie S1 and Table S1). These results suggest that affinity-optimizing SNVs can contribute to migration defects during heart development.

The ETS3 and ETS4 SNVs both cause a 3-fold increase in affinity, creating sites with 0.34 and 0.52 affinities respectively. Surprisingly, migration defects are seen in 9.9% of embryos containing the FoxF ETS3 T to G enhancer driving FoxF despite the fact this mutation only creates a 0.34 affinity site. The FoxF ETS4 T to C driving FoxF causes migration defects in 8.0% of embryos (Figure S5 and Table S1). These results indicate that affinity-optimizing changes of ≥3-fold, even those that result in relatively low-affinity sites, can alter cell behavior. Furthermore, the disproportionate impact of the affinity-optimizing SNV in ETS3, which is located near two other important sites (Figure S3), suggests that enhancer grammar may influence the functional consequences of affinity-optimizing SNVs within genomes. Having seen the impact of each SNV individually on TVC migration, we next wanted to see what impact all three affinity-optimizing variants would have on migration. However, embryos with the FoxF 3 SNVs driving FoxF expression do not survive to tailbud stage and are deformed beyond recognition. We therefore could not assess the number of migration defects in ATMs as we could not identify these populations of cells.

### Affinity-optimizing SNVs disrupt heart development in the most severe case leading to two beating hearts

To determine the impact of migration defects on heart development, we followed 52 FoxF ETS1 T to C embryos with migration defects from the larval stage to the juvenile stage. Strikingly, 79% of embryos with migration defects had abnormally developed hearts (Figure 4). Phenotypes ranged from enlarged hearts to two distinct beating hearts found within the same pericardium (Movie **S3**). As the animals age, hearts of *Ciona* with the optimizing SNV could not pump blood cells (Movie S4-8) and these animals began dying at twelve days after metamorphosis. Thus, single base-pair changes that cause a ≥3-fold increase in the affinity of a single ETS site contribute to changes in gene expression and changes in heart development.

### Low-affinity ETS sites are prevalent within *Ciona* developmental heart enhancers

Having seen the importance of low-affinity ETS sites within the FoxF enhancer, we wanted to see if low-affinity binding sites were found within other *Ciona* developmental heart enhancers. Using published ATAC-seq data for *Ciona* heart cells at 6.5hr post fertilization, a time at which FGF signaling is received by the heart precursor cells, we identified 15,174 putative *Ciona* heart enhancers (Racioppi et al., 2019). The mean affinity of putative ETS sites within these putative enhancers was 0.12, suggesting that low-affinity sites are prevalent within *Ciona* putative heart enhancers. We next wanted to see how many of these putative enhancers have an ETS site that could have a SNV that increases the affinity of ETS sites by ≥ 3-fold. We found 6,618 (44%) of *Ciona* putative developmental heart enhancers contain at least one ETS site in which a SNV could increase the affinity by ≥ 3-fold (Figure 5B). This indicates that use of low-affinity binding sites is prevalent within *Ciona* developmental heart enhancers and that these enhancers are vulnerable to variants that increase the affinity of ETS sites.

### Mouse and human GATA4 enhancers contain suboptimal affinity ETS sites

The role of FGF signaling in specification and migration of heart cells is conserved from flies to vertebrates (Harvey, 2002; Zaffran and Frasch, 2002). To see if the use of low-affinity ETS sites within developmental heart enhancers is also conserved to vertebrates we first looked at a well-studied and functionally characterized ETS-dependent vertebrate developmental heart enhancer, the GATA4 G9 enhancer (Schachterle et al., 2012). This enhancer has been tested via reporter assay in mouse and drives expression in the endocardial and myocardial layers of the heart from e8.5 to e11.5 (Schachterle et al., 2012). Ablating four of these ETS sites leads to loss of expression, suggesting that these ETS sites are required for expression (Schachterle et al., 2012). While these sites have been annotated, the affinity of these sites has not been studied. We measured the affinity of all ten ETS sites using mouse ETS-1 PBM data. Strikingly, all ten ETS sites are very low-affinity ranging from 0.09 to 0.12 relative to consensus. The four sites that were previously ablated causing loss of expression have affinities of 0.11, 0.12, 0.10, and 0.12 respectively. These findings demonstrate that very low-affinity ETS are necessary for heart-specific expression from the mouse GATA4 G9 enhancer. The mouse GATA4 G9 enhancer is highly conserved within the human genome (Schachterle et al., 2012). Indeed, the human homologue of the mouse GATA4 G9 enhancer also contains very low-affinity binding sites that are almost identical to those found within the mouse GATA4 G9 enhancer (Figure 5A). This demonstrates on a functionally validated enhancer that very low-affinity binding sites are required for heart-specific expression in mice and that this is likely conserved to humans.

### Human putative developmental heart enhancers contain low-affinity ETS binding sites, and the majority of these enhancers contain an ETS site that can be optimized by a SNV

Having seen that the GATA4 G9 enhancer in vertebrates contains low-affinity sites important for enhancer activity, we next conducted a genome-wide analysis to identify putative human developmental heart enhancers. We annotated putative developmental heart enhancers using epigenomic datasets derived from primary fetal human hearts and iPSC-derived embryonic-like cardiovascular progenitor cells at timepoints when FGF signaling is important for heart development (Sylva et al., 2014). This collection of epigenomic datasets contains ATAC-seq and ChIP-seq for p300, Gata4, Tbx5, Med1. The samples, time points, and references of these datasets are detailed in Table S2. From this data, we identified 197,963 putative human developmental heart enhancers. These putative developmental heart enhancers contain very low-affinity ETS sites with a mean relative affinity of 0.12, which mirrors the mean ETS affinity seen in *Ciona* heart developmental enhancers (Figure 5B). Strikingly, 66% of these human putative developmental heart enhancers contain an ETS site that, upon a single base pair change, has a ≥ 3-fold increase in affinity (Figure 5B).

### Affinity-optimizing SNVs are associated with cardiac traits

We next wanted to investigate if affinity-optimizing variants are associated with cardiac traits in humans. To achieve this, we obtained from the UK Biobank 368,241 non-coding GWAS variants (p<1e-4) for cardiac traits: pulse rate, pulse pressure, and QRS duration. We chose a p<1e-4 threshold as it is becoming increasingly common to use values below the genome-wide significance threshold (p<5e-8) to identify putative variants that contribute to cardiac traits, especially when used in combination with other functional features such as epigenomic marks, or transcription factor binding sites (Wang et al., 2016). We identified 446 GWAS ETS affinity-optimizing SNVs using the subthreshold p<1e-4, of which 15% (67/446) are found within developmental heart enhancers. Considering only variants that reach genome-wide significance (p<5e-8), we find 125 GWAS ETS affinity optimizing SNVs (MAF 0.01-0.49, Table S3), 20% (22/125) of these occur within mapped developmental heart enhancers. As we do not yet have an exhaustive map of all putative regulatory elements, some of the non-coding GWAS ETS affinity-optimizing SNVs not overlapping could lie within currently unmapped heart enhancers. The molecular underpinnings of very few cardiac trait and disease GWAS associations are known. The potential functional impact of these 446 GWAS ETS affinity-optimizing SNVs make them excellent candidates for causality.

### Affinity-optimizing SNV associated with QRS duration occurs within an SCN5A enhancer

One of the GWAS variants, rs12635900, is an affinity-optimizing SNV which leads to an 8-fold increase in ETS binding affinity (Figure 6A). This variant is associated with QRS duration and is within a putative developmental heart enhancer, as marked by H3K27Ac, that interacts with the SCN5A promoter (Man et al., 2019; Richter et al., 2020). SCN5A is the major cardiac sodium channel which plays an important role in excitation and contraction of the heart (Li et al., 2018; Veerman et al., 2015). Deletions, and gain- or loss-of-function mutations in SCN5A are associated with a spectrum of human cardiac conduction traits and diseases including QRS duration, PR interval, bradycardia, atrial fibrillation, and Brugada syndrome (Daimi et al., 2019; Li et al., 2018). Precise expression of SCN5A in cardiomyocytes is required for normal heart conduction, and mice with higher levels of SCN5A show changes in heart conduction (Liu et al., 2015; Zhang et al., 2007). The putative SCN5A fetal heart enhancer contains a cluster of low-affinity sites ranging from 0.09 – 0.25 relative binding affinity (Figure 6B). The presence of low-affinity ETS sites within this putative enhancer is consistent with our previous analysis in *Ciona*, mouse and human cardiac enhancers. SCN5A is expressed in cardiomyocytes and thus we wanted to validate that this putative enhancer region drives expression in human cardiomyocytes. We predicted that the affinity-optimizing variant could lead to increased enhancer activity in human cardiomyocytes.

### The SCN5A affinity-optimizing SNV causes gain-of-function expression in human iPSC-derived cardiomyocytes

To test the reference SCN5A putative enhancer and the enhancer harboring the affinity-optimizing SNV, we used human iPSC-CMs (D’Antonio-Chronowska et al., 2019; Panopoulos et al., 2017). We transfected the reference enhancer and the enhancer containing the affinity-optimizing SNV driving GFP into human iPSC-CMs. Samples were collected 96 hours after transfection and GFP expression from both the reference and affinity-optimizing SNV enhancer were quantified by qPCR. The affinity-optimizing SNV leads to a statistically significant 1.8-fold increase in expression (p = 0.0016) (Figure 6). Thus, in human developmental heart enhancers, affinity-optimizing variants drive gain-of-function gene expression consistent with our findings in *Ciona*. Given that this affinity-optimizing SNV is a common variant associated with QRS duration, and that this enhancer interacts with the SCN5A promoter (Man et al., 2019), our results suggests that the affinity-optimizing variant may increase SCN5A expression and contribute to changes in QRS duration. Overexpression of SCN5A in mice is known to alter heart conduction phenotypes (Liu et al., 2015; Zhang et al., 2007). This analysis provides proof-of-principle that low-affinity sites could contribute to cardiac phenotypes in humans. Nine other significant GWAS SNVs are within this same SCN5A putative enhancer, highlighting the importance of overlaying mechanistic understanding to predict causal variants. Furthermore, our approach provides a framework to integrate searches for affinity-optimizing SNVs, epigenomic, interaction data and GWAS analysis to pinpoint causal enhancer variants within a sea of linked variants.

## Discussion

There are millions of non-coding variants associated with disease states, complex traits, evolutionary adaptations and phenotypic variation (Maurano et al., 2012; Tak and Farnham, 2015; Visel et al., 2009). However, pinpointing these causal variants is a huge challenge as they are typically embedded within a sea of inert variants (Gallagher and Chen-Plotkin, 2018). It is therefore critical that we develop generalizable approaches to pinpoint causal enhancer variants. Here we use a mechanistic understanding of the rules governing enhancer activity to pinpoint causal enhancer variants involved in heart development and cardiac traits. We find that suboptimal or low-affinity binding sites are prevalent within developmental heart enhancers in *Ciona*, mouse and humans. This creates a pervasive vulnerability within genomes whereby single nucleotide changes can dramatically increase affinity, leading to gain-of-function gene expression that contributes to phenotypic changes in both the *Ciona* and human heart. Our mechanistic approach provides a much-needed functionally validated and systematic framework that works across different enhancers, cell types and species to pinpoint causal enhancer variants contributing to enhanceropathies, phenotypic diversity and evolutionary changes.

### Suboptimal-affinity ETS sites are a prevalent feature of chordate developmental heart enhancers

Our genome-wide analysis finds that low-affinity ETS sites are a common feature of developmental heart enhancers in chordates. The use of low-affinity sites within developmental heart enhancers across *Ciona*, mouse and human is highlighted by the sites found in the FoxF, GATA4 and SCN5A enhancers which range from 0.10-0.25. Ablation of these low-affinity sites both within *Ciona* (sites of 0.11 to 0.24 affinity) and in mouse (sites of 0.10 to 0.12 affinity) leads to a dramatic reduction in enhancer activity demonstrating that these low-affinity sites are functional. Our findings highlight the importance and vastly underappreciated contribution of very low-affinity binding sites to regulation of gene expression across organisms. The use of low-affinity sites is also important within enhancers activated by other transcription factors that act downstream of signaling pathways and pleiotropic factors (Crocker et al., 2015, 2016; Farley et al., 2015; Rowan et al., 2010; Tsai et al., 2017). Thus, the use of low-affinity binding sites to encode precise expression is likely a generalizable principle that applies to developmental enhancers and beyond.

### Affinity-optimizing SNVs cause gain-of-function enhancer activity which can contribute to phenotypes

A major consequence of using such low-affinity sites within developmental enhancers is that single base-pair changes can dramatically optimize the affinity of a site, causing gain-of-function expression and phenotypic changes within the heart. Our genome-wide analysis indicates that the majority of putative developmental heart enhancers in both *Ciona* and humans harbor low-affinity ETS sites that can be optimized by a SNV. In *Ciona*, functional validation shows that affinity-optimizing SNVs lead to loss of tissue-specific expression and migration defects that alter heart development. In humans, we identify 446 GWAS ETS affinity-optimizing SNVs significantly associated with cardiac traits. One of these GWAS ETS affinity-optimizing SNVs, associated with QRS duration, occurs within an SCN5A enhancer and leads to increased expression relative to the reference enhancer. Higher levels of SCN5A in mouse cause changes in heart conduction (Liu et al., 2015; Zhang et al., 2007). Given the prevalence of low-affinity binding sites within enhancers and the ease with which these can be optimized, affinity-optimizing variants could be a common and underappreciated mechanism driving phenotypic change in both disease and evolutionary contexts.

### Enhancer grammar likely modifies the impact of affinity-optimizing SNVs

In this study, we identify three different ETS affinity-optimizing variants within the FoxF enhancer, each of which leads to gain-of-function enhancer activity and migration defects. However, optimizing different ETS sites within the enhancer contributes to phenotypes in a way that is nonlinear with affinity. The ETS3 site, is the lowest affinity site within the FoxF enhancer, yet it seems to be particularly important. Ablation of this site causes the greatest reduction in expression. While increasing the affinity of this site to 0.34, a 3-fold increase in binding affinity, leads to almost the same number of migration defects as the SNV within ETS1 that leads to an 8-fold increase to a final affinity of 0.97. Similarly, ETS4 has a greater impact on phenotype than expected based on the 3-fold increase in affinity. The proximity of the ETS3 and ETS4 sites to other important binding sites (ATTA and Ebox) may be boosting the impact of the affinity-optimizing SNVs. These results indicate that the affinity, along with the relative context of sites within the enhancer, or the enhancer grammar, impacts expression and the consequence of any optimizing variant (Jindal and Farley, 2021). Future studies to understand the interplay between affinity-optimization and enhancer grammar could further refine our ability to predict causal enhancer variants.

### Affinity-optimizing SNVs may contribute to evolution of multi-chambered hearts

While we have mainly focused on the disease implications of affinity-optimizing SNVs, there is a fine line between creation of novel phenotypes that are beneficial or detrimental (Tournamille et al., 1995). The *Ciona* heart like all invertebrate chordates has a single chambered heart, while all extant vertebrates have at least two chambers. The evolution of a dual chambered heart in vertebrates is thought to involve recruitment of additional precursor cells to the ventral midline (Davidson et al., 2006). Our study indicates that affinity-optimizing variants could contribute to more cells migrating to the ventral midline and creation of another compartment within the pericardium, thus a multichambered heart. It is possible that some animals with these multichambered hearts could gradually be modified to exploit the selective advantage of an independent inflow and outflow tract. Previous studies have found that overexpressing ETS in the ATMs can lead to the same phenotype (Davidson et al., 2006). Our study and the previous studies suggest that changes both at the level of FGF signaling and within enhancers responsive to FGF could contribute to evolution of a multi-chambered heart.

### Predicting causal enhancer variants by layering mechanistic understanding with other genomic datasets

Approaches to pinpoint causal enhancer variants include overlaying epigenomic and interaction data with GWAS and eQTL data (Cano-Gamez and Trynka, 2020; Mahajan et al., 2018). However, even with these approaches, there are often many variants within an enhancer associated with a particular trait. For example, we use epigenomic data from human fetal heart to find an SCN5A putative enhancer and 4-C interaction data to link this to the SCN5A promoter (Man et al., 2019; Richter et al., 2020). Within this SCN5A enhancer, we find nine SNVs with similar p-values all associated with QRS duration. We pinpointed the one variant that causes a dramatic increase in ETS affinity and find that this variant drives higher levels of expression (Figure 6D). Thus, using a mechanistic approach on top of the already available datasets helps pinpoint enhancer variants that cause gain-of-function gene expression.

### Affinity-optimizing variants lead to gain-of-function gene expression in diverse enhancers involved in different cell types and developmental processes

The FoxF and SCN5A enhancers are vastly different; one is important for migration of heart cells in *Ciona* (Beh et al., 2007), while the other is important for heart conduction. The FoxF enhancer is located within 3kb of its target promoter (Beh et al., 2007), while SCN5A enhancer is located over 100Mb away from its target promoter (Man et al., 2019). In our accompanying paper, we find affinity-optimizing variants within the ZRS enhancer cause ectopic expression of Shh in the developing mouse limb bud and polydactyly. These enhancers in Ciona, mouse and human are all very different and yet they are all governed by the use of low-affinity ETS sites to encode precise patterns of gene expression. Violating this regulatory principle in all of these settings leads to loss of precise expression. Within the context of the FoxF and ZRS enhancers, this is the loss of tissuespecificity. While in the human SCN5A enhancer, it is the loss of a particular level of expression. These studies collectively demonstrate that affinity-optimizing variants can lead to gain-of-function gene expression that can contribute to phenotypic changes in a variety of diverse organisms and tissues.

### Searching for affinity-optimizing variants is a systematic, generalizable strategy to pinpoint causal enhancer variants across species and cell types

Changes in binding affinity have been linked to changes in gene expression and phenotypes in a handful of studies (Bond et al., 2004; French et al., 2013; Grant et al., 1996; Huang et al., 2014; Jia et al., 2009; Tuupanen et al., 2009). Notably, these are one off examples rather than systematic approaches to find causal variants across disparate enhancers, associated with different diseases and traits across species and cell types. Collectively, our studies demonstrate that searching for affinity-optimizing variants is a generalizable mechanistic approach that can pinpoint causal variants across a variety of diverse enhancers across cell types and species. As this approach relies on a mechanistic understanding of enhancers, it has the power to be equally effective at identifying causal variants regardless of their frequency or origin (i.e., inherited, somatic, common, rare, private). The biological context, and other factors such as selection will likely alter the effect size of variants, therefore making it easier to validate certain variants within the endogenous locus, such as rare variants of strong effect size found within the ZRS.

### Violations in regulatory principles governing enhancers can pinpoint causal enhancer variants

Given the importance of FGF signaling for heart development and function, the prevalence of low-affinity sites within cardiac enhancers, and the ease with which these sites can be optimized, we suggest ETS affinity-optimizing single nucleotide variants likely contribute to congenital heart disease and cardiac traits. The ubiquitous role of FGF in other developmental programs (Jindal et al., 2015; Xie et al., 2020) and cancers (Grose and Dickson, 2005) indicates that ETS affinity-optimizing single nucleotide variants are likely also involved in other enhanceropathies.

Finding rules that generalize across different enhancers relies on understanding the regulatory principles that govern enhancer function. One such regulatory principle is the use of suboptimal-affinity binding sites to encode tissue-specific expression and control levels of expression. Here we show that SNVs that violate this regulatory principle can cause gain-of-function gene expression and phenotypic changes. Identification of other regulatory principles and subsequently variants that violate these could provide further approaches to pinpoint causal enhancer variants at scale.

## Acknowledgements

We thank the Farley and Frazer labs, especially Matteo D’Antonio for helpful discussions. We thank Jim Posakony and Mike Levine for a critical reading of the manuscript.

## Author contributions

E.K.F, G.A.J, A.T.B, K.A.F, A.D-C. designed experiments. G.A.J., A.T.B., J.L.G., F.L., S.H.L., A.D-C. conducted experiments. J.J.S, R.O.L., A.K. conducted bioinformatic analyses; E.K.F wrote the manuscript. All authors were involved in editing the manuscript.

## Methods

### Key resource table

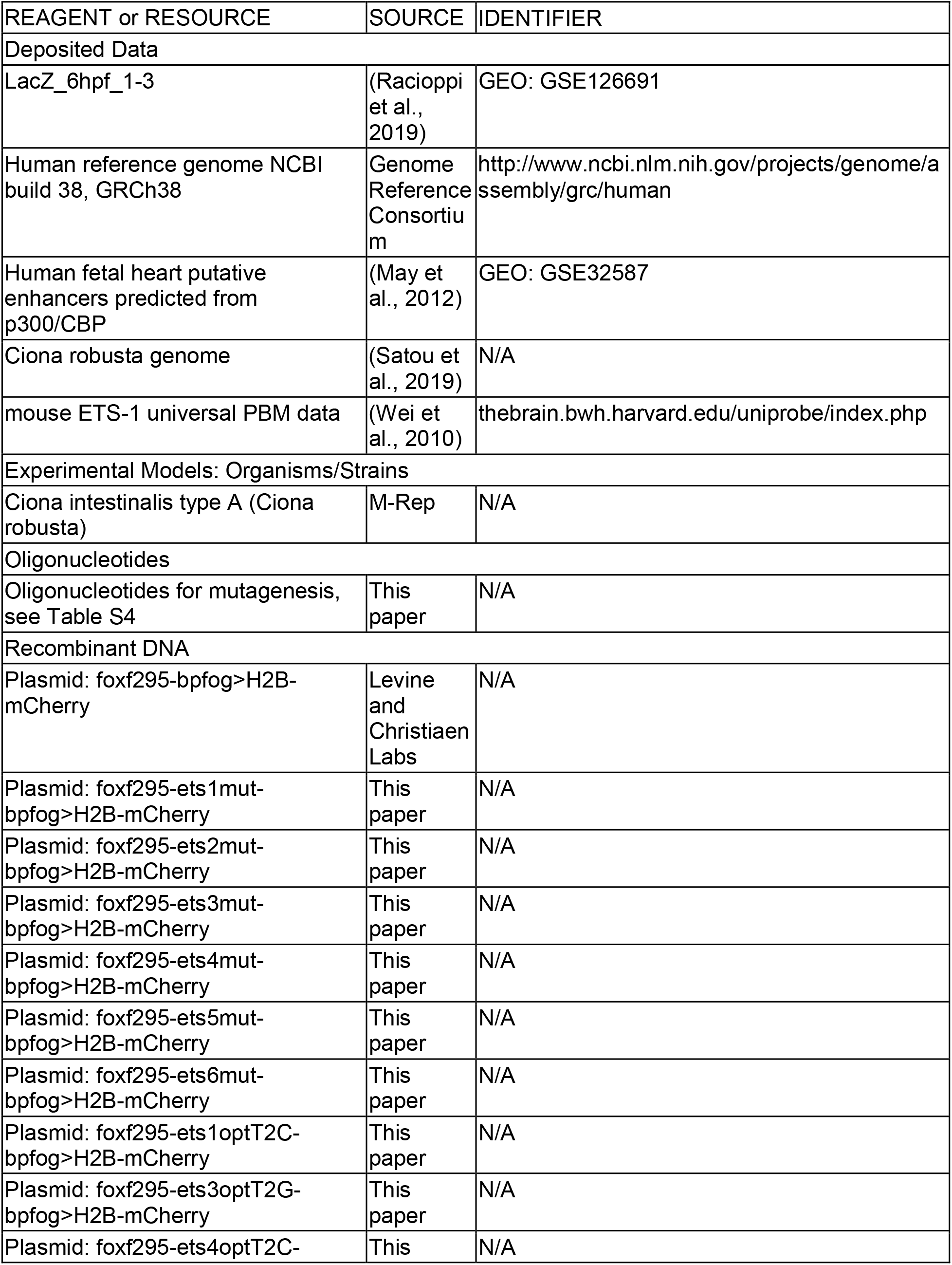
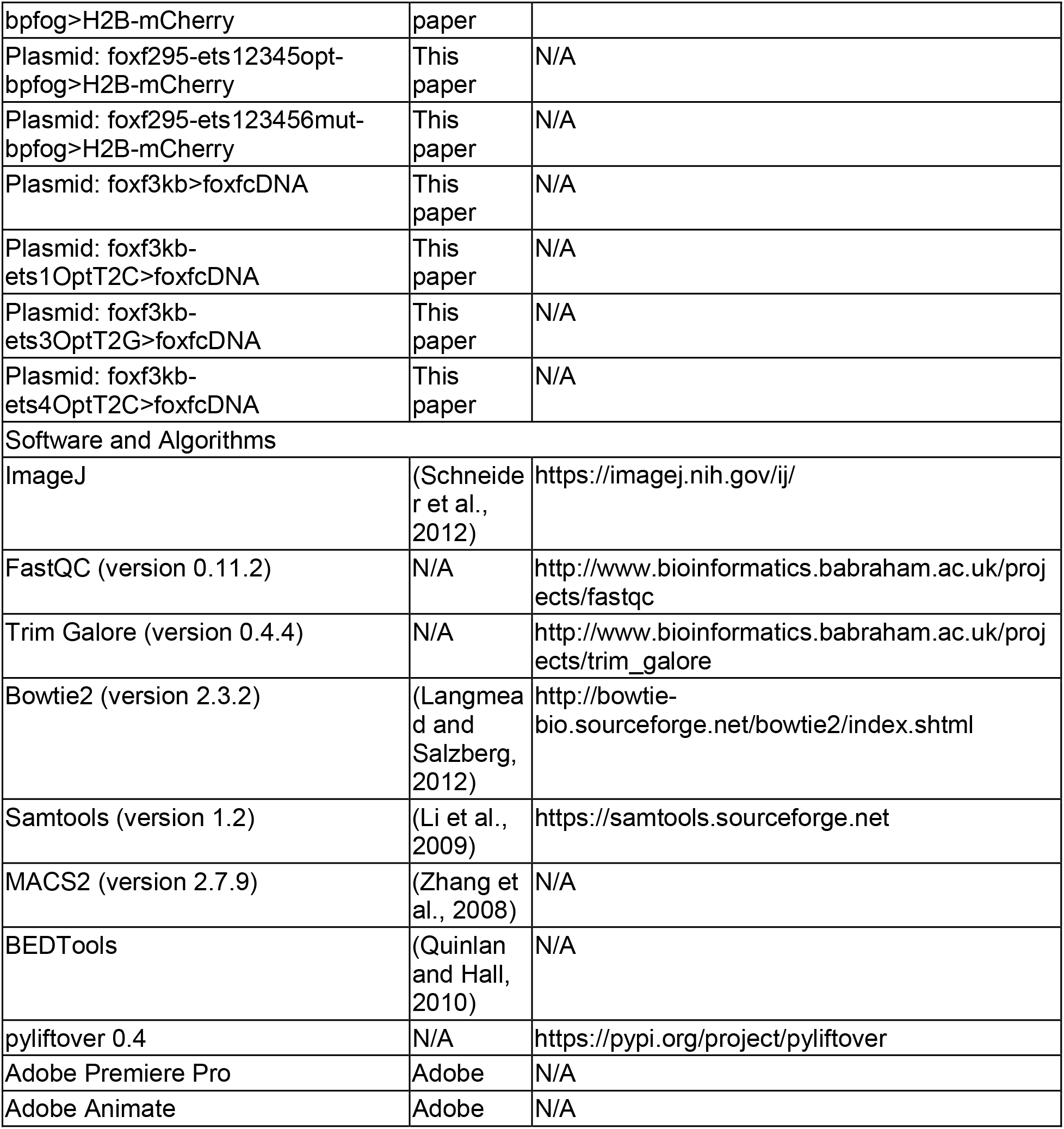

## RESOURCE AVAILABILITY

### Lead contact

Further information and requests for resources and reagents should be directed to and will be fulfilled by the lead contact, Emma Farley.

### Materials availability

Plasmids generated in this study will be deposited in Addgene by the date of publication.

### Data and code availability

- Microscopy and scoring data reported in this paper will be shared by the lead contact upon request.
- All original code **will be** deposited at GitHub (https://github.com/jsolvason/cell2022-heart) and **will be** made publicly available as of the date of publication. DOIs **will be** listed in the key resources table.
- Any additional information required to reanalyze the data reported in this paper is available from the lead contact upon request.

#### Data and materials availability

- Putative Ciona developmental heart enhancers: We will upload to GEO the bed we created using FastQ files from Racciopi et al., 2019 (GEO accession GSE126691).
- Putative human developmental heart enhancers: We provide the references for all these datasets in Supplementary table 2. We can also provide all regions as a supplementary bed file.

## EXPERIMENTAL MODEL AND SUBJECT DETAILS

### Tunicates

Adult *C. intestinalis* type A aka *Ciona robusta* (obtained from M-Rep) were maintained under constant illumination in seawater (obtained from Reliant Aquariums) at 18 C. Ciona are hermaphroditic, therefore there is only one possible sex for individuals. Age or developmental stage of the embryos studied are indicated in the main text.

## METHOD DETAILS

### Electroporation

Dechorionation, in vitro fertilization and electroporation were performed as described previously in (Christiaen et al., 2009). 70 μg DNA was resuspended in 100 μL water and added to 400 μL of 0.96 M D-mannitol. Typically for each electroporation, eggs and sperm were collected from 6 adults. Embryos were fixed at the appropriate developmental stage for 15 minutes in 3.7% formaldehyde. The tissue was then cleared in a series of washes of 0.3% Triton-X in PBS and then of 0.01% Triton-X in PBS. Samples were mounted in Prolong Gold. Differential interference contrast microscopy was used to obtain transmitted light micrographs with an Olympus FV3000, using the 40X objective. The same microscope was used to obtain all mCherry images. All constructs were electroporated with three biological replicates, unless otherwise noted.

### Counting embryos

For each experiment, slides were counted blind. In each experiment, all comparative constructs were present. Fifty embryos were counted for each biological replicate, unless otherwise noted.

### Mutagenesis and cloning

The FoxF enhancer construct was a gift from the Levine and Christiaen labs. The construct consists of the following elements in order: AscI, FoxF enhancer, XbaI, bpfog promoter, NotI, H2B-mCherry, EcoRI. Point mutations were introduced into the construct using mutagenesis with partially overlapping primers with 3’-overhangs.To clone the construct with all 6 sites mutated from GGAW to GCAW, we (1) ordered the foxf enhancer synthesized in two parts from IDT, (2) amplified each part using one forward and one reverse primer, (3) cut each with a Type IIS restriction enzyme SapI, (4) ligated together using ligase, and (5) finally cloned the resulting fragment into the FoxF enhancer construct using the AscI and XbaI sites. The constructs driving foxf cDNA expression were cloned using a more complicated strategy, since foxf cDNA contains 1 NotI and 2 EcoRI sites. Foxf cDNA with the NotI site mutated away with a silent mutation was cloned into the foxf enhancer construct using the NotI and MfeI restriction sites. This scheme works because MfeI and EcoRI have the same 4 base pair overhang. Foxf cDNA with the NotI site mutated away with a silent mutation was synthesized by Twist Bioscience. The cDNA sequence was found in Aniseed with the Gene Model ID KH2012:KH.C3.170. The SCN5A variant fragments were synthesized by Twist Bioscience and ordered with Esp3I restriction sites at each end. The variant fragments were then cloned into the vector to test in iPSC-CMs with Esp3I. The vector consists of the following elements in order: SCN5A fragment, Supercore promoter, unc76-EGFP, bGH-polyA.

### Acquisition of Images

For enhancers being compared, images were taken from electroporations performed on the same day using identical settings. Two exposure times were taken for each construct. For representative images, embryos were chosen that represented the average from counting data. All images are subsequently cropped to an appropriate size. In each figure, the same exposure time for each image is shown to allow direct comparison, which occasionally leads to overexposed images being used for stronger constructs (e.g., Fig 1C).

### Counting Larvae for Migration Defects

Embryos were transferred from gelatin-coated electroporation plates to 15cm uncoated plastic plates filled with filtered seawater at 7 hpf. All well-developed larvae on the plate were counted from 16-20 hpf and scored for presence or absence of normal ATM development. All larvae with migration defects were marked for heart morphology analysis in the juvenile stage. For each replicate 10-20 larva with normal ATM development were selected for heart morphology analysis in the juvenile stage.

### Juvenile Heart Morphology Analysis and Imaging

Juvenile hearts were analyzed for morphological defects starting at 48hrs after metamorphosis. Hearts with abnormal structure or function were scored as “deformed”. Images and video of hearts were taken on an Axiozoom microscope with a mounted iPhone 7. Video editing was done in Adobe Premiere Pro. Heartbeat animations were made in Adobe Animate.

### TVC Migration Time-lapse Imaging

Embryos were electroporated with MesP 2kb>GFP as previously described and checked for GFP fluorescence at 6 hpf. Well-developed embryos with strong GFP expression were transferred to a glass-bottom microscopy plate in seawater. Z-stacks were taken of each embryo every 30 minutes for 8 hours with an Olympus FV3000 microscope with a 20x objective lens. Resulting time-lapses were analyzed in Fiji. A max projection was taken of the GFP channel and merged with the DIC. Time-lapses were cropped and rotated in Adobe Photoshop.

### Scoring Relative Affinities

Relative affinities were calculated using high-throughput binding datasets (Hume et al., 2015) (thebrain.bwh.harvard.edu/uniprobe/index.php). They were calculated using median signal intensities of mouse ETS-1 universal protein binding microarray data from the UniProbe database (Wei et al., 2010). It has previously been shown that the binding specificity of ETS-1 across 600 million years from flies to humans is conserved and thus the use of mouse ETS-1 to calculate ETS affinity is a valid approach (Nitta et al., 2015). The percentage of relative affinities represent the fold changes of median signal intensities of the native 8-mer motifs compared with the optimal 8-mer motif for ETS.

### ATAC-seq Data Analysis

#### Alignment of ATAC-seq reads

ATAC-seq alignment was performed following the methods in (Racioppi et al., 2019). Raw reads from 6 hours post fertilization (hpf) (GEO accession GSE126691, LacZ_6hpf_1-3, LacZ_10hpf_1-4) were first preprocessed by FastQC (version 0.11.2, http://www.bioinformatics.babraham.ac.uk/projects/fastqc). Adaptors were trimmed using Trim Galore (version 0.4.4, http://www.bioinformatics.babraham.ac.uk/projects/trim_galore) and trimmed reads were aligned to the Ciona robusta genome (Satou et al., 2019) using Bowtie2 (version 2.3.2, (Langmead and Salzberg, 2012)) with the parameters –very-sensitive -p 16 -X 1000. Reads with mapping quality score > 30 were kept for downstream analysis using SAMtools (version 1.2, (Li et al., 2009)). Mitochondrial reads were removed using bash command egrep -v. Following examination of read quality of the 6hpf data, replicate 2 was removed. Read pileup (BAM) files for replicates 1 and 3 at 6hpf were merged using the SAMtools merge function and peaks were called using MACS2 (version 2.7.9) (Zhang et al., 2008) (--nomodel --bdg --g 99000000 -f BAMPE -q 0.01). To correct for nonspecific sequencing biases, we subtracted gDNA from these libraries (Toenhake et al., 2018).

### Calculating top ATAC peaks

To find the highest confidence peaks in our dataset, we calculated the area under the curve (AUC) as the sum of the read depth at each position in the peak. Bedtools genomecov (-bga) and intersect functions were used to calculate the read depth at every genomic position within all the called peaks (Quinlan and Hall, 2010). Peaks with AUCs in the top 90% were kept and the rest were discarded.

#### Downloading putative enhancers

Human fetal heart putative enhancers were identified from epigenomic datasets from Table S2. Any coordinates for hg18 or hg19 were lifted over to hg38 using pyliftover 0.4 (https://pypi.org/project/pyliftover).

#### Risk of optimization of ETS contained in putative developmental heart enhancers

We first determined the distribution of ETS affinities in putative developmental heart enhancers. We collected affinities for all ETS sites defined as NNGGAWNN. Once all of the ETS binding site affinities were determined, we then performed all possible point mutations to each binding site and plotted those which resulted in ≥3-fold change. We reported the number of putative heart enhancers which contained ≥1 ETS which could be affinity-optimized ≥3 fold by a point mutation.

#### Searching disease-associated mutations which optimize ETS site

We searched for ETS affinity-optimizing mutations in UK Biobank GWAS for 3 cardiac traits: Pulse Rate, QRS Duration, Pulse Pressure. All coordinates were lifted over to hg38 before analysis using pyliftover. We filtered for non-coding bedtools intersect -v to remove exonic GWAS variants using UCSC refGene exons. We considered a variant to be in an ETS site if it resided in the NNGGAWNN ETS site at any N or W position. For variants occurring in the W position, we only allowed for A>T and T>A mutations as this maintains the IUPAC definition for W. We defined the variant as being ETS optimizing if the alt allele increased the affinity ≥3-fold. Allele frequency (AF) data was taken from UK Biobank. The minor allele frequency is defined as the frequency of the minor allele, meaning if the UK Biobank provided the AF for the major allele, we calculated the minor AF as 1-AF.

### iPSC-CM transfection and collection

iPSC-CMs from line UDID_104 (D’Antonio-Chronowska et al., 2019) were thawed and seeded onto gelatin-coasted 48-well plates at a density of 1.3 x 10^5^/cm^2^. The iPSC-CMs were transfected with 325nM/well of plasmid containing either the reference enhancer or the enhancer harboring the affinity-optimizing SNV driving GFP using ViaFect™ Transfection Reagent (Promega) at a 6:1 ratio in a medium containing 10% FBS and 5uM (Y27632) ROCK Inhibitor (Sigma). 16hrs post transfection and every other day, medium was changed. Cells were then collected 96hrs post transfection and stored in TriZol reagent frozen at −80°C. Transfection for each allele was performed in duplicate.

### RNA extraction, cDNA synthesis, qPCR

Total RNA was extracted using TriZol extraction and treated to remove contaminating DNA (TURBO DNA-free DNase digestion kit, Ambion). Subsequent mRNA was isolated (Dynabeads mRNA Isolation kit, Invitrogen) and was used for cDNA synthesis (Transcriptor High Fidelity cDNA Synthesis kit, Sigma-Aldrich). GFP expression levels were quantified with qPCR using three different reference genes (RPS29, HMBS, SCN5A). The qPCR was run with iQ SYBR Green Supermix (Bio-rad) with anneal temperature of 59°C (see table S4 for primers).

## QUANTIFICATION AND STATISTICAL ANALYSIS

For comparisons of counting data, chi-squared test was used in Excel with the CHISQ.TEST function. qPCR expression data was analyzed using the delta-delta-Ct method in Excel and statistically analyzed using the Student’s T-test in Excel.

## Supplemental Figures

**Figure S1.**
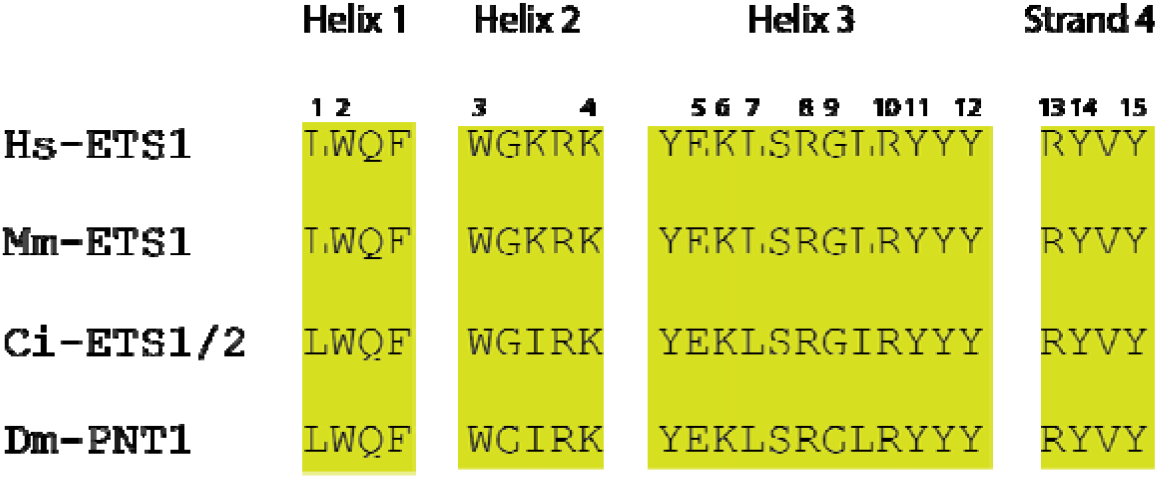
ETS binding domains conservation. ETS domains showing amino acid residues that contact specific nucleotides, based on published crystal structures of ETS domain DNA complexes from human, mouse, *Ciona*, and *Drosophila* (Farley et al., 2015). Amino acid residues contacting DNA are numbered from 1 to 15.

**Figure S2.**
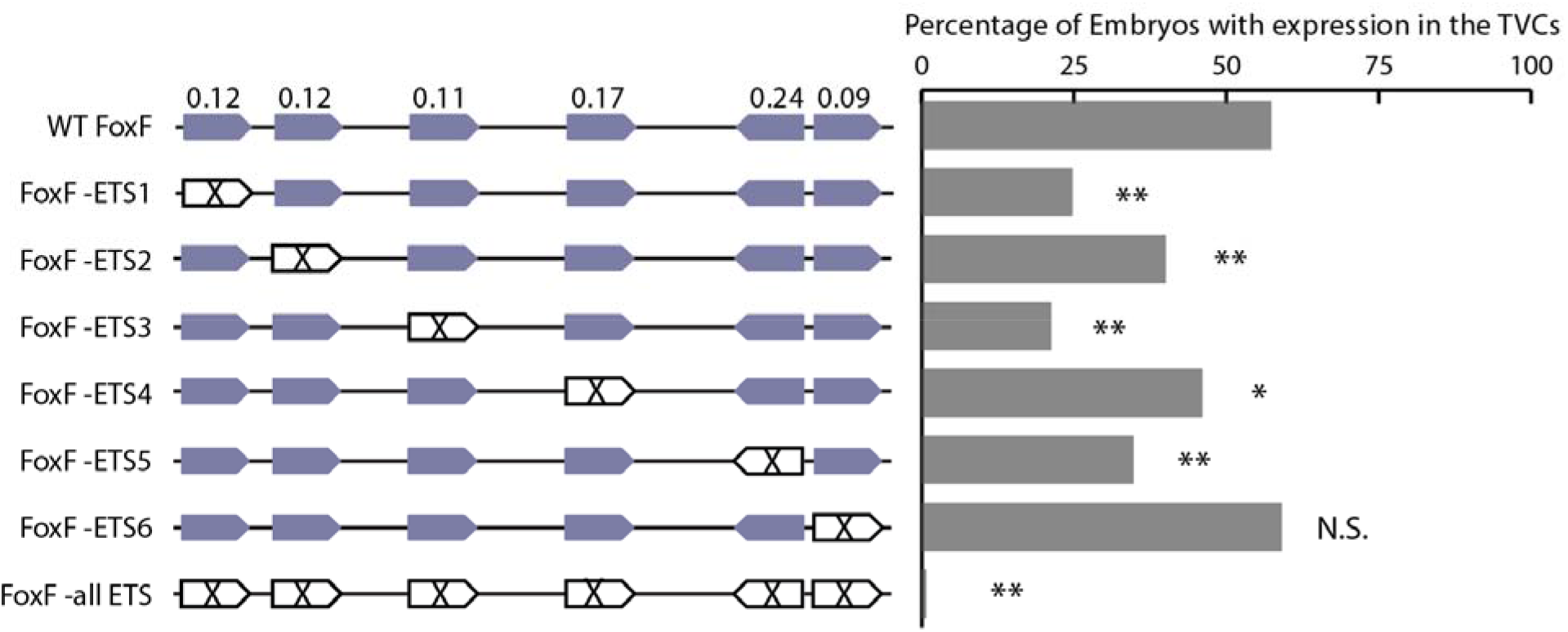
Counting data for deletion of ETS sites within FoxF enhancer. Schematics show the enhancers that were electroporated into embryos. Diagrams indicate the WT FoxF fragment and X indicates GGAW > GCAW mutation. Graph shows the proportions of embryos and levels of expression in TVCs (3 replicates, n ≥ 50 embryos for each replicate). P values are obtained by using Chi-square test comparing each condition to WT FoxF enhancer: ** P < 0.005, * P <0.05, N.S. not significant.

**Figure S3.**
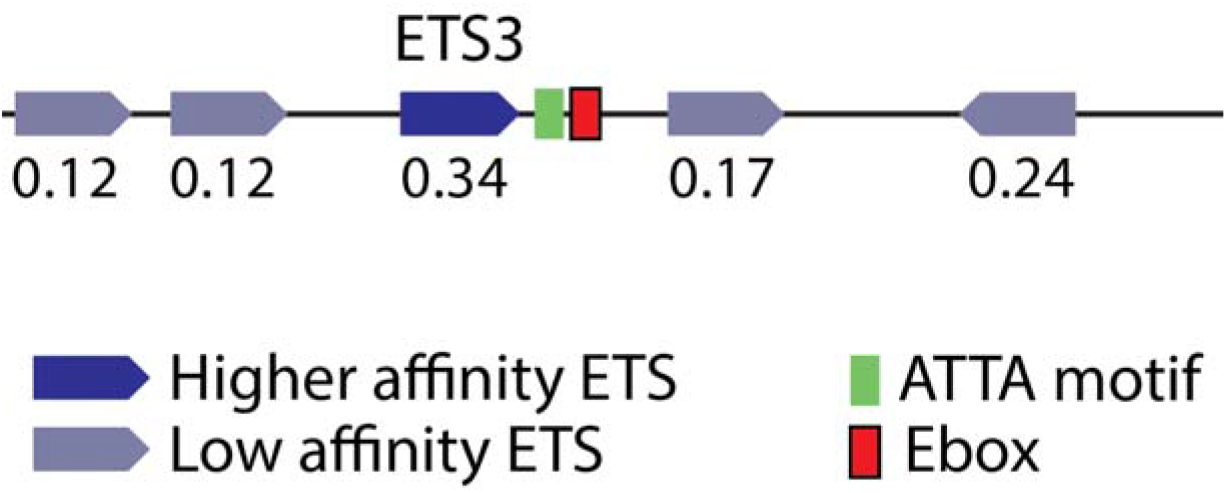
Other sites within the FoxF enhancer. Schematic of FoxF enhancer with ETS3 SNV, increasing the affinity from 0.11 to 0.34. ATTA motif and Ebox binding site are shown directly upstream of ETS3. Ebox is the putative binding site for Mesp, a major cardiac transcription factor (Bondue and Blanpain, 2010).

**Figure S4.**
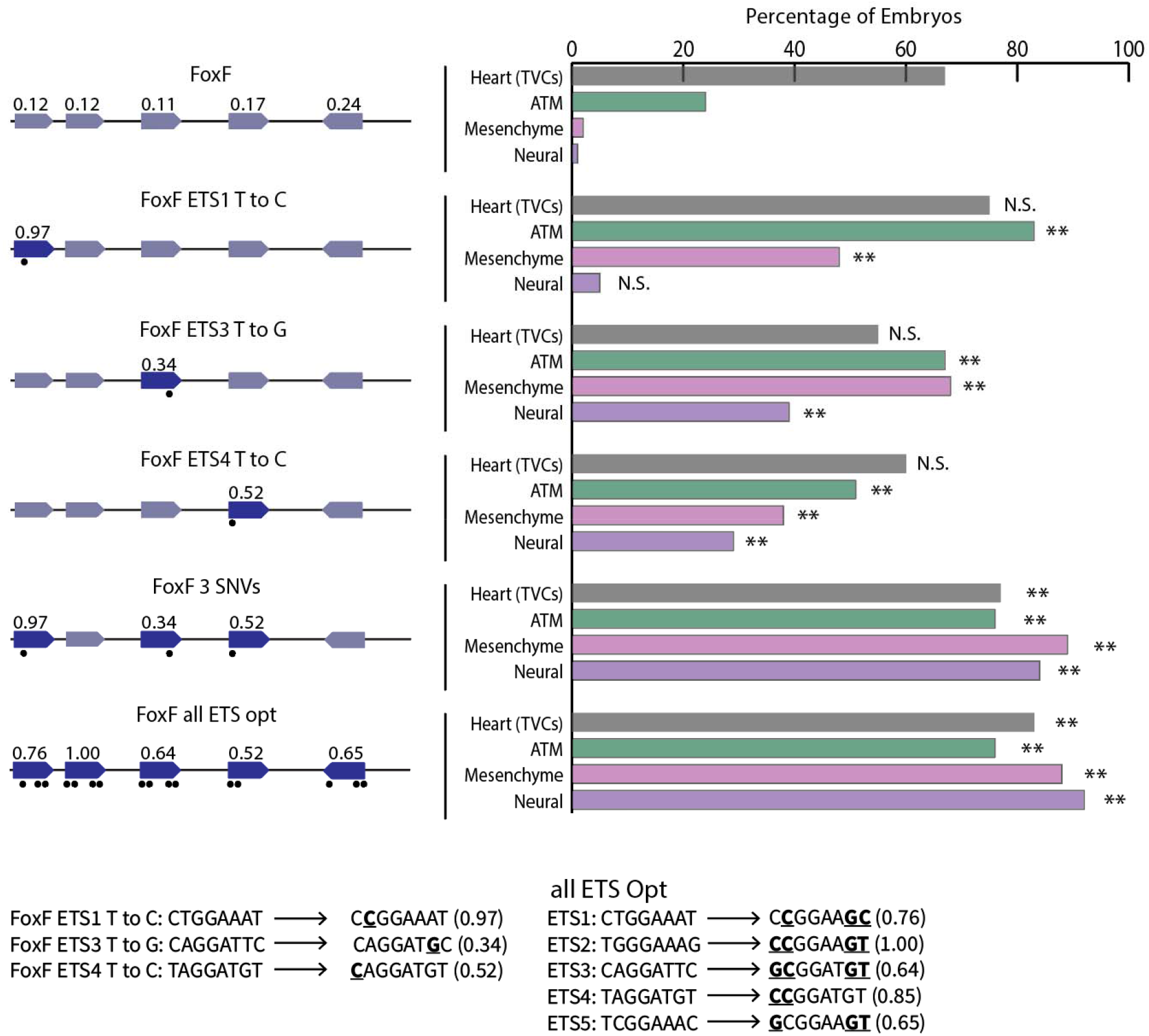
Counting data for affinity-optimizing single base-pair changes. Scoring for embryos electroporated with the WT FoxF enhancer, FoxF SNVs, FoxF 3 SNVs, and the FoxF all ETS opt enhancer (3 replicates, n = 50 embryos for each replicate). Schematics show the enhancers that were electroporated into embryos, values are affinities, dots show the nucleotide changes (3 replicates, n ≥ 38 embryos for each replicate). P values are obtained by using Chisquare test comparing each condition to the WT FoxF enhancer: ** P < 0.005, * P <0.05, N.S. not significant. The sequences of each site before and after the nucleotide change is shown. Abbreviated tissue is Anterior tail muscle (ATM).

**Figure S5.**
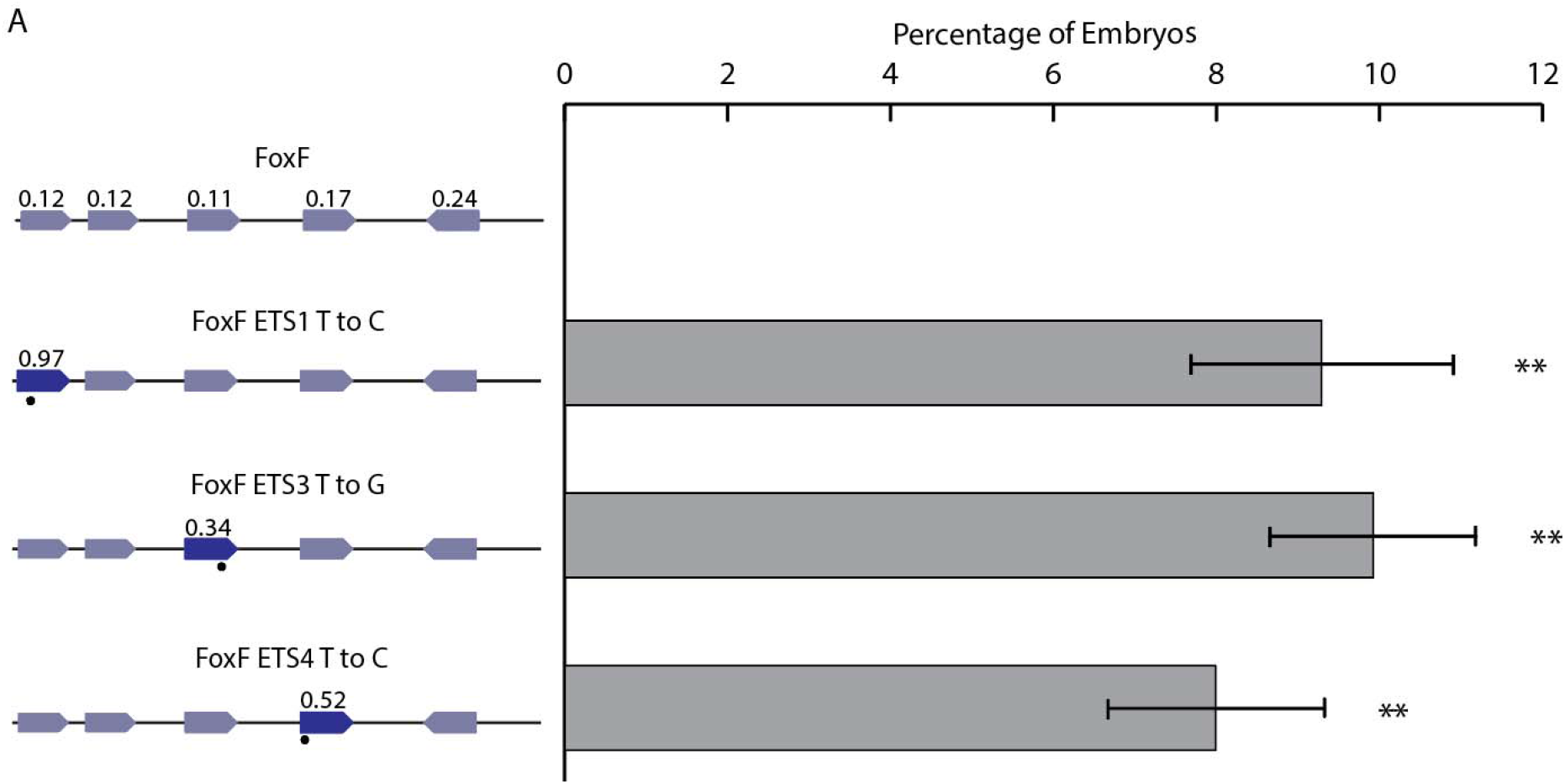
Counting of migration defects in tailbud embryos with affinity-optimizing mutations. **A.** We scored the migration of ATMs into the ventral midline in between 250 and 380 embryos per condition. The WT FoxF enhancer driving FoxF drove no ATM migration in embryos (0/286). FoxF ETS1 T to C, FoxF ETS3 T to G and FoxF ETS4 T to C driving FoxF caused ATM migration defects in 9.3% (n=23/250), 9.9% (n=37/364) and 8.0% (n=31/380) of embryos respectively. P values are obtained by using Chi-square test comparing to WT FoxF enhancer: ** P < 0.005. The single base-pair changes made are shown by the dots in the enhancer schematic.

**Table S1:**
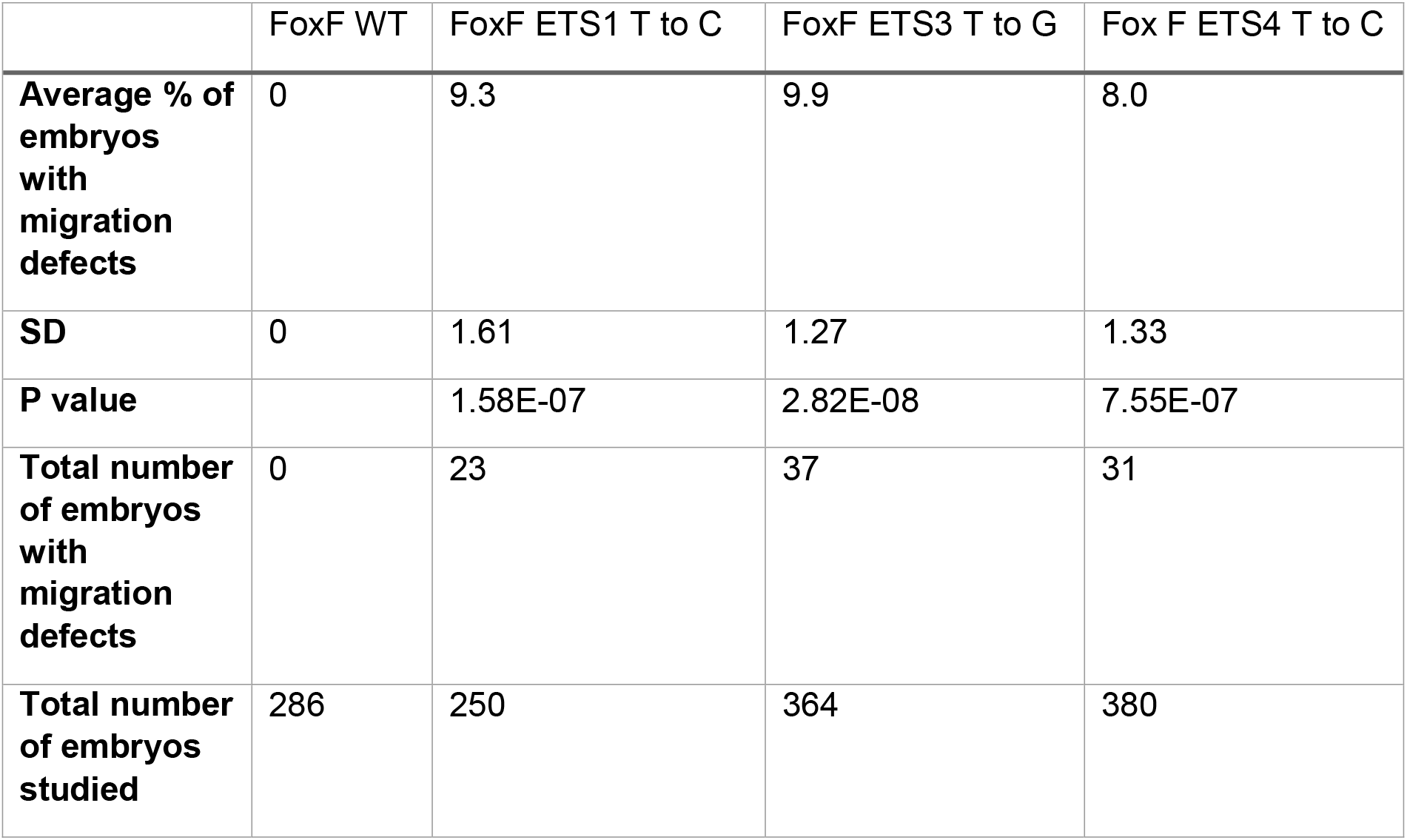
Numbers for counting of migration defects in tailbud embryos with affinity-optimizing mutations. Three replicates were carried out for FoxF ETS1 T to C and WT. Two replicates of FoxF ETS1 T to G and FoxF ETS4 T to C. P values are obtained using Chi-squared test comparing to the WT FoxF enhancer.

**Table S2:** Epigenomic datasets used to identify human putative fetal heart enhancers. The table provides information on all samples, timepoints, epigenomic approach and references to the publication used to define putative developmental heart enhancers.

| Sample | Time Point | Sequencing Method | Reference |
| --- | --- | --- | --- |
| IPSC-Cardio Myocyte | Day 15,25 | ChIP H3K27ac | (Benaglio et al., 2019) |
| IPSC-Cardio Myocyte | Day 15,25 | ATAC-seq | (Benaglio et al., 2019) |
| IPSC-Cardiac Mesoderm | Day 8 | ATAC-seq | (Richter et al., 2020) |
| IPSC-Cardiac Mesoderm | Day 17 | ATAC-seq | (Richter et al., 2020) |
| IPSC-CVPC | Day 25 | ATAC-seq | (D'Antonio et al., 2021) |
| IPSC-CVPC | Day 25 | ChIP H3K27ac | (D'Antonio et al., 2021) |
| Fetal Human Heart Biopsy | Week 8-17 | ChIP H3K27ac | (Richter et al., 2020) |
| Fetal Human Heart Biopsy | Week 16 | ChIP p300 | (May et al., 2012) |
| IPSC-CPC | Day 7 | ATAC-seq | (Ang et al., 2016) |
| IPSC-CPC | Day 7 | ChIP H3K27ac | (Ang et al., 2016) |
| IPSC-CPC | Day 7 | ChIP GATA4 | (Ang et al., 2016) |
| IPSC-CPC | Day 7 | ChIP Tbx5 | (Ang et al., 2016) |
| IPSC-CPC | Day 7 | ChIP Med1 | (Ang et al., 2016) |

**Table S3: GWAS ETS-Affinity Optimizing SNVs associated with cardiac traits: Pulse pressure, pulse rate and QRS duration.** The table includes the location of the SNV, the affinity, the affinity fold change and the minor allele frequency.

**Table S4: Oligonucleotides used in this study.**

**Movie S1. Ectopic FoxF expression in ATMs drives cell migration.** (Top) Embryo with WT FoxF enhancer driving the FoxF gene shows normal TVC migration to the ventral midline while ATMs remain in the tail and begin to elongate. (Bottom) Embryo electroporated with FoxF ETS1 T to C enhancer driving FoxF gene shows migration of TVCs and ATMs together to the ventral midline. Timelapses taken from late neurula to late tailbud stages (St. 16-23).

**Movie S2. Juvenile *Ciona* with normal TVC migration develop proper heart morphology**. The juvenile *Ciona* heart is highly simplistic, consisting of a single linear chamber within the heart cavity in which peristaltic motion cycles blood through the body.

**Movie S3. Juvenile *Ciona* with ATM migration develop an extra heart.** Embryos with FoxF ETS1 T to C driving FoxF exhibiting migrated ATMs were tracked through metamorphosis. This juvenile developed extra heart tissue which contracts independently of the main heart tube.

**Movie S4. Juvenile *Ciona* hearts with normal morphology function to cycle blood through the body.** Same movie as S2 but with blood cells labelled. Blood is able to move freely through the heart tube in a functioning juvenile heart.

**Movie S5. Juvenile *Ciona* hearts with an expanded phenotype are often unable to properly function.** Embryos with FoxF ETS1 T to C driving FoxF exhibiting migration defects were tracked through metamorphosis. This embryo has extra chambers and abnormal morphology, the peristaltic motion is disrupted, and blood is unable to be circulated.

**Movie S6. An improperly functioning heart is highly detrimental to organism health and viability.** When the blood cannot circulate, it becomes trapped and clotted, eventually leading to death of the organism. Whole animal blood tracking showing little movement of blood throughout the body. (Same individual as movie S5, 6 days older).

**Movie S7. An improperly functioning heart is highly detrimental to organism health and viability.** Same animal as movie S6 zoomed in on blood tracking in heart. Very little blood movement and severe clotting present.

**Movie S8. An improperly functioning heart is highly detrimental to organism health and viability.** Same animal as in movie S6 but longer video, to show the lack of blood flow over a longer time frame. This movie has no blood cell tracking.

## References

Ang, Y.-S., Rivas, R.N., Ribeiro, A.J.S., Srivas, R., Rivera, J., Stone, N.R., Pratt, K., Mohamed, T.M.A., Fu, J.-D., Spencer, C.I., et al. (2016). Disease Model of GATA4 Mutation Reveals Transcription Factor Cooperativity in Human Cardiogenesis. Cell 167, 1734–1749.e22. https://doi.org/10.1016/j.cell.2016.11.033.

Beh, J., Shi, W., Levine, M., Davidson, B., and Christiaen, L. (2007). FoxF is essential for FGF-induced migration of heart progenitor cells in the ascidian Ciona intestinalis. Development 134, 3297–3305. https://doi.org/10.1242/dev.010140.

Beiman, M., Shilo, B.Z., and Volk, T. (1996). Heartless, a Drosophila FGF receptor homolog, is essential for cell migration and establishment of several mesodermal lineages. Genes Dev. 10, 2993–3002. https://doi.org/10.1101/gad.10.23.2993.

Benaglio, P., D’Antonio-Chronowska, A., Ma, W., Yang, F., Young Greenwald, W.W., Donovan, M.K.R., DeBoever, C., Li, H., Drees, F., Singhal, S., et al. (2019). Allele-specific NKX2-5 binding underlies multiple genetic associations with human electrocardiographic traits. Nat. Genet. 51, 1506–1517. https://doi.org/10.1038/s41588-019-0499-3.

Berger, M.F., and Bulyk, M.L. (2009). Universal protein-binding microarrays for the comprehensive characterization of the DNA-binding specificities of transcription factors. Nat. Protoc. 4, 393–411. https://doi.org/10.1038/nprot.2008.195.

Bond, G.L., Hu, W., Bond, E.E., Robins, H., Lutzker, S.G., Arva, N.C., Bargonetti, J., Bartel, F., Taubert, H., Wuerl, P., et al. (2004). A single nucleotide polymorphism in the MDM2 promoter attenuates the p53 tumor suppressor pathway and accelerates tumor formation in humans. Cell 119, 591–602. https://doi.org/10.1016/j.cell.2004.11.022.

Bondue, A., and Blanpain, C. (2010). Mesp1: a key regulator of cardiovascular lineage commitment. Circ. Res. 107, 1414–1427. https://doi.org/10.1161/CIRCRESAHA.110.227058.

Boyle, E.A., Li, Y.I., and Pritchard, J.K. (2017). An Expanded View of Complex Traits: From Polygenic to Omnigenic. Cell 169, 1177–1186. https://doi.org/10.1016/j.cell.2017.05.038.

Buckingham, M., Meilhac, S., and Zaffran, S. (2005). Building the mammalian heart from two sources of myocardial cells. Nat. Rev. Genet. 6, 826–835. https://doi.org/10.1038/nrg1710.

Cano-Gamez, E., and Trynka, G. (2020). From GWAS to Function: Using Functional Genomics to Identify the Mechanisms Underlying Complex Diseases. Front. Genet. 11.

Christiaen, L., Davidson, B., Kawashima, T., Powell, W., Nolla, H., Vranizan, K., and Levine, M. (2008). The Transcription/Migration Interface in Heart Precursors of Ciona intestinalis. Science 320, 1349–1352. https://doi.org/10.1126/science.1158170.

Christiaen, L., Wagner, E., Shi, W., and Levine, M. (2009). Electroporation of Transgenic DNAs in the Sea Squirt Ciona. Cold Spring Harb. Protoc. 2009, pdb.prot5345. https://doi.org/10.1101/pdb.prot5345.

Cooley, J., Whitaker, S., Sweeney, S., Fraser, S., and Davidson, B. (2011). Cytoskeletal polarity mediates localized induction of the heart progenitor lineage. Nat. Cell Biol. 13, 952–957. https://doi.org/10.1038/ncb2291.

Cota, C.D., and Davidson, B. (2015). Mitotic Membrane Turnover Coordinates Differential Induction of the Heart Progenitor Lineage. Dev. Cell 34, 505–519. https://doi.org/10.1016/j.devcel.2015.07.001.

Crocker, J., Abe, N., Rinaldi, L., McGregor, A.P., Frankel, N., Wang, S., Alsawadi, A., Valenti, P., Plaza, S., Payre, F., et al. (2015). Low Affinity Binding Site Clusters Confer Hox Specificity and Regulatory Robustness. Cell 160, 191–203. https://doi.org/10.1016/j.cell.2014.11.041.

Crocker, J., Preger-Ben Noon, E., and Stern, D.L. (2016). Chapter Twenty-Seven - The Soft Touch: Low-Affinity Transcription Factor Binding Sites in Development and Evolution. In Current Topics in Developmental Biology, P.M. Wassarman, ed. (Academic Press), pp. 455–469.

Daimi, H., Khelil, A.H., Neji, A., Ben Hamda, K., Maaoui, S., Aranega, A., Be Chibani, J., and Franco, D. (2019). Role of SCN5A coding and non-coding sequences in Brugada syndrome onset: What’s behind the scenes? Biomed. J. 42, 252–260. https://doi.org/10.1016/j.bj.2019.03.003.

D’Antonio, M., Arthur, T.D., Nguyen, J.P., Matsui, H., D’Antonio-Chronowska, A., and Frazer, K.A. (2021). Fine mapping spatiotemporal mechanisms of genetic variants underlying cardiac traits and disease. BioRxiv https://doi.org/10.1101/2021.09.01.458619.

D’Antonio-Chronowska, A., Donovan, M.K.R., Young Greenwald, W.W., Nguyen, J.P., Fujita, K., Hashem, S., Matsui, H., Soncin, F., Parast, M., Ward, M.C., et al. (2019). Association of Human iPSC Gene Signatures and X Chromosome Dosage with Two Distinct Cardiac Differentiation Trajectories. Stem Cell Rep. 13, 924–938. https://doi.org/10.1016/j.stemcr.2019.09.011.

Davidson, B. (2007). Ciona intestinalis as a model for cardiac development. Semin. Cell Dev. Biol. 18, 16–26. https://doi.org/10.1016/j.semcdb.2006.12.007.

Davidson, B., Shi, W., Beh, J., Christiaen, L., and Levine, M. (2006). FGF signaling delineates the cardiac progenitor field in the simple chordate, Ciona intestinalis. Genes Dev. 20, 2728–2738. https://doi.org/10.1101/gad.1467706.

Delker, R.K., Ranade, V., Loker, R., Voutev, R., and Mann, R.S. (2019). Low affinity binding sites in an activating CRM mediate negative autoregulation of the Drosophila Hox gene Ultrabithorax. PLoS Genet. 15, e1008444. https://doi.org/10.1371/journal.pgen.1008444.

Delsuc, F., Brinkmann, H., Chourrout, D., and Philippe, H. (2006). Tunicates and not cephalochordates are the closest living relatives of vertebrates. Nature 439, 965–968. https://doi.org/10.1038/nature04336.

Farley, E.K., Olson, K.M., Zhang, W., Brandt, A.J., Rokhsar, D.S., and Levine, M.S. (2015). Suboptimization of developmental enhancers. Science 350, 325–328. https://doi.org/10.1126/science.aac6948.

Farley, E.K., Olson, K.M., Zhang, W., Rokhsar, D.S., and Levine, M.S. (2016). Syntax compensates for poor binding sites to encode tissue specificity of developmental enhancers. Proc. Natl. Acad. Sci. 113, 6508–6513. https://doi.org/10.1073/pnas.1605085113.

Fisher, R.A. (1918). The correlation between relatives on the supposition of Mendelian inheritance. Trans R Soc Edinb 52, 399–433.

Frazer, K.A., Murray, S.S., Schork, N.J., and Topol, E.J. (2009). Human genetic variation and its contribution to complex traits. Nat. Rev. Genet. 10, 241–251. https://doi.org/10.1038/nrg2554.

French, J.D., Ghoussaini, M., Edwards, S.L., Meyer, K.B., Michailidou, K., Ahmed, S., Khan, S., Maranian, M.J., O’Reilly, M., Hillman, K.M., et al. (2013). Functional variants at the 11q13 risk locus for breast cancer regulate cyclin D1 expression through long-range enhancers. Am. J. Hum. Genet. 92, 489–503. https://doi.org/10.1016/j.ajhg.2013.01.002.

Fuqua, T., Jordan, J., van Breugel, M.E., Halavatyi, A., Tischer, C., Polidoro, P., Abe, N., Tsai, A., Mann, R.S., Stern, D.L., et al. (2020). Dense and pleiotropic regulatory information in a developmental enhancer. Nature 587, 235–239. https://doi.org/10.1038/s41586-020-2816-5.

Gallagher, M.D., and Chen-Plotkin, A.S. (2018). The Post-GWAS Era: From Association to Function. Am. J. Hum. Genet. 102, 717–730. https://doi.org/10.1016/j.ajhg.2018.04.002.

Grant, S.F., Reid, D.M., Blake, G., Herd, R., Fogelman, I., and Ralston, S.H. (1996). Reduced bone density and osteoporosis associated with a polymorphic Sp1 binding site in the collagen type I alpha 1 gene. Nat. Genet. 14, 203–205. https://doi.org/10.1038/ng1096-203.

Grose, R., and Dickson, C. (2005). Fibroblast growth factor signaling in tumorigenesis. Cytokine Growth Factor Rev. 16, 179–186. https://doi.org/10.1016/j.cytogfr.2005.01.003.

Harvey, R.P. (2002). Patterning the vertebrate heart. Nat. Rev. Genet. 3, 544–556. https://doi.org/10.1038/nrg843.

House, S.L., House, B.E., Glascock, B., Kimball, T., Nusayr, E., Schultz, E.J., and Doetschman, T. (2010). Fibroblast Growth Factor 2 Mediates Isoproterenol-induced Cardiac Hypertrophy through Activation of the Extracellular Regulated Kinase. Mol Cell Pharmacol 2, 143–154.

Huang, Q., Whitington, T., Gao, P., Lindberg, J.F., Yang, Y., Sun, J., Väisänen, M.-R., Szulkin, R., Annala, M., Yan, J., et al. (2014). A prostate cancer susceptibility allele at 6q22 increases RFX6 expression by modulating HOXB13 chromatin binding. Nat. Genet. 46, 126–135. https://doi.org/10.1038/ng.2862.

Jia, L., Landan, G., Pomerantz, M., Jaschek, R., Herman, P., Reich, D., Yan, C., Khalid, O., Kantoff, P., Oh, W., et al. (2009). Functional enhancers at the gene-poor 8q24 cancer-linked locus. PLoS Genet. 5, e1000597. https://doi.org/10.1371/journal.pgen.1000597.

Jindal, G.A., and Farley, E.K. (2021). Enhancer grammar in development, evolution, and disease: dependencies and interplay. Dev. Cell 56, 575–587. https://doi.org/10.1016/j.devcel.2021.02.016.

Jindal, G.A., Goyal, Y., Burdine, R.D., Rauen, K.A., and Shvartsman, S.Y. (2015). RASopathies: unraveling mechanisms with animal models. Dis. Model. Mech. 8, 769–782. https://doi.org/10.1242/dmm.020339.

Kribelbauer, J.F., Rastogi, C., Bussemaker, H.J., and Mann, R.S. (2019). Low-Affinity Binding Sites and the Transcription Factor Specificity Paradox in Eukaryotes. Annu. Rev. Cell Dev. Biol. 35, 357–379. https://doi.org/10.1146/annurev-cellbio-100617-062719.

Langmead, B., and Salzberg, S.L. (2012). Fast gapped-read alignment with Bowtie 2. Nat. Methods 9, 357–359. https://doi.org/10.1038/nmeth.1923.

Lettice, L.A., Hill, A.E., Devenney, P.S., and Hill, R.E. (2008). Point mutations in a distant sonic hedgehog cis-regulator generate a variable regulatory output responsible for preaxial polydactyly. Hum. Mol. Genet. 17, 978–985. https://doi.org/10.1093/hmg/ddm370.

Lettice, L.A., Williamson, I., Wiltshire, J.H., Peluso, S., Devenney, P.S., Hill, A.E., Essafi, A., Hagman, J., Mort, R., and Grimes, G. (2012). Opposing functions of the ETS factor family define Shh spatial expression in limb buds and underlie polydactyly. Dev. Cell 22, 459–467.

Levine, M. (2010). Transcriptional Enhancers in Animal Development and Evolution. Curr. Biol. 20, R754–R763. https://doi.org/10.1016/j.cub.2010.06.070.

Li, H., Handsaker, B., Wysoker, A., Fennell, T., Ruan, J., Homer, N., Marth, G., Abecasis, G., Durbin, R., and 1000 Genome Project Data Processing Subgroup (2009). The Sequence Alignment/Map format and SAMtools. Bioinformatics 25, 2078–2079. https://doi.org/10.1093/bioinformatics/btp352.

Li, W., Yin, L., Shen, C., Hu, K., Ge, J., and Sun, A. (2018). SCN5A Variants: Association With Cardiac Disorders. Front. Physiol. 9, 1372. https://doi.org/10.3389/fphys.2018.01372.

Liu, G.X., Remme, C.A., Boukens, B.J., Belardinelli, L., and Rajamani, S. (2015). Overexpression of SCN5A in mouse heart mimics human syndrome of enhanced atrioventricular nodal conduction. Heart Rhythm 12, 1036–1045. https://doi.org/10.1016/j.hrthm.2015.01.029.

Mahajan, A., Taliun, D., Thurner, M., Robertson, N.R., Torres, J.M., Rayner, N.W., Payne, A.J., Steinthorsdottir, V., Scott, R.A., Grarup, N., et al. (2018). Fine-mapping type 2 diabetes loci to single-variant resolution using high-density imputation and islet-specific epigenome maps. Nat. Genet. 50, 1505–1513. https://doi.org/10.1038/s41588-018-0241-6.

Man, J.C.K., Mohan, R.A., Boogaard, M. van den, Hilvering, C.R.E., Jenkins, C., Wakker, V., Bianchi, V., Laat, W. de, Barnett, P., Boukens, B.J., et al. (2019). An enhancer cluster controls gene activity and topology of the SCN5A-SCN10A locus in vivo. Nat. Commun. 10, 4943. https://doi.org/10.1038/s41467-019-12856-5.

Mathieson, I. (2021). The omnigenic model and polygenic prediction of complex traits. Am. J. Hum. Genet. 108, 1558–1563. https://doi.org/10.1016/j.ajhg.2021.07.003.

Maurano, M.T., Humbert, R., Rynes, E., Thurman, R.E., Haugen, E., Wang, H., Reynolds, A.P., Sandstrom, R., Qu, H., Brody, J., et al. (2012). Systematic Localization of Common Disease-Associated Variation in Regulatory DNA. Science 337, 1190–1195. https://doi.org/10.1126/science.1222794.

May, D., Blow, M.J., Kaplan, T., McCulley, D.J., Jensen, B.C., Akiyama, J.A., Holt, A., Plajzer-Frick, I., Shoukry, M., Wright, C., et al. (2012). Large-scale discovery of enhancers from human heart tissue. Nat. Genet. 44, 89–93. https://doi.org/10.1038/ng.1006.

Nitta, K.R., Jolma, A., Yin, Y., Morgunova, E., Kivioja, T., Akhtar, J., Hens, K., Toivonen, J., Deplancke, B., Furlong, E.E.M., et al. (2015). Conservation of transcription factor binding specificities across 600 million years of bilateria evolution. ELife 4, e04837. https://doi.org/10.7554/eLife.04837.

Panopoulos, A.D., D’Antonio, M., Benaglio, P., Williams, R., Hashem, S.I., Schuldt, B.M., DeBoever, C., Arias, A.D., Garcia, M., Nelson, B.C., et al. (2017). iPSCORE: A Resource of 222 iPSC Lines Enabling Functional Characterization of Genetic Variation across a Variety of Cell Types. Stem Cell Rep. 8, 1086–1100. https://doi.org/10.1016/j.stemcr.2017.03.012.

Quinlan, A.R., and Hall, I.M. (2010). BEDTools: a flexible suite of utilities for comparing genomic features. Bioinformatics 26, 841–842. https://doi.org/10.1093/bioinformatics/btq033.

Racioppi, C., Wiechecki, K.A., and Christiaen, L. (2019). Combinatorial chromatin dynamics foster accurate cardiopharyngeal fate choices. ELife 8, e49921. https://doi.org/10.7554/eLife.49921.

Reifers, F., Walsh, E.C., Leger, S., Stainier, D.Y.R., and Brand, M. (2000). Induction and differentiation of the zebrafish heart requires fibroblast growth factor 8 (fgf8/acerebellar). Development 127, 225–235.

Richter, F., Morton, S.U., Kim, S.W., Kitaygorodsky, A., Wasson, L.K., Chen, K.M., Zhou, J., Qi, H., Patel, N., DePalma, S.R., et al. (2020). Genomic analyses implicate noncoding de novo variants in congenital heart disease. Nat. Genet. 52, 769–777. https://doi.org/10.1038/s41588-020-0652-z.

Rowan, S., Siggers, T., Lachke, S.A., Yue, Y., Bulyk, M.L., and Maas, R.L. (2010). Precise temporal control of the eye regulatory gene Pax6 via enhancer-binding site affinity. Genes Dev. 24, 980–985. https://doi.org/10.1101/gad.1890410.

Satou, Y., Nakamura, R., Yu, D., Yoshida, R., Hamada, M., Fujie, M., Hisata, K., Takeda, H., and Satoh, N. (2019). A Nearly Complete Genome of Ciona intestinalis Type A (C. robusta) Reveals the Contribution of Inversion to Chromosomal Evolution in the Genus Ciona. Genome Biol. Evol. 11, 3144–3157. https://doi.org/10.1093/gbe/evz228.

Schachterle, W., Rojas, A., Xu, S.-M., and Black, B.L. (2012). ETS-dependent regulation of a distal Gata4 cardiac enhancer. Dev. Biol. 361, 439–449. https://doi.org/10.1016/j.ydbio.2011.10.023.

Schneider, C.A., Rasband, W.S., and Eliceiri, K.W. (2012). NIH Image to ImageJ: 25 years of image analysis. Nat. Methods 9, 671–675. https://doi.org/10.1038/nmeth.2089.

Sharrocks, A.D. (2001). The ETS-domain transcription factor family. Nat. Rev. Mol. Cell Biol. 2, 827–837. https://doi.org/10.1038/35099076.

Smemo, S., Tena, J.J., Kim, K.-H., Gamazon, E.R., Sakabe, N.J., Gómez-Marín, C., Aneas, I., Credidio, F.L., Sobreira, D.R., Wasserman, N.F., et al. (2014). Obesity-associated variants within FTO form long-range functional connections with IRX3. Nature 507, 371–375. https://doi.org/10.1038/nature13138.

Sotoodehnia, N., Isaacs, A., de Bakker, P.I.W., Dörr, M., Newton-Cheh, C., Nolte, I.M., van der Harst, P., Müller, M., Eijgelsheim, M., Alonso, A., et al. (2010). Common variants in 22 loci are associated with QRS duration and cardiac ventricular conduction. Nat. Genet. 42, 1068–1076. https://doi.org/10.1038/ng.716.

Spivakov, M. (2014). Spurious transcription factor binding: non-functional or genetically redundant? BioEssays News Rev. Mol. Cell. Dev. Biol. 36, 798–806. https://doi.org/10.1002/bies.201400036.

Swanson, C.I., Schwimmer, D.B., and Barolo, S. (2011). Rapid Evolutionary Rewiring of a Structurally Constrained Eye Enhancer. Curr. Biol. 21, 1186–1196. https://doi.org/10.1016/j.cub.2011.05.056.

Swenson, B.R., Louie, T., Lin, H.J., Méndez-Giráldez, R., Below, J.E., Laurie, C.C., Kerr, K.F., Highland, H., Thornton, T.A., Ryckman, K.K., et al. (2019). GWAS of QRS duration identifies new loci specific to Hispanic/Latino populations. PloS One 14, e0217796. https://doi.org/10.1371/journal.pone.0217796.

Sylva, M., van den Hoff, M.J.B., and Moorman, A.F.M. (2014). Development of the human heart. Am. J. Med. Genet. A. 164A, 1347–1371. https://doi.org/10.1002/ajmg.a.35896.

Tak, Y.G., and Farnham, P.J. (2015). Making sense of GWAS: using epigenomics and genome engineering to understand the functional relevance of SNPs in non-coding regions of the human genome. Epigenetics Chromatin 8, 57. https://doi.org/10.1186/s13072-015-0050-4.

Toenhake, C.G., Fraschka, S.A.-K., Vijayabaskar, M.S., Westhead, D.R., van Heeringen, S.J., and Bártfai, R. (2018). Chromatin Accessibility-Based Characterization of the Gene Regulatory Network Underlying Plasmodium falciparum Blood-Stage Development. Cell Host Microbe 23, 557–569.e9. https://doi.org/10.1016/j.chom.2018.03.007.

Tournamille, C., Colin, Y., Cartron, J.P., and Kim, C.L.V. (1995). Disruption of a GATA motif in the Duffy gene promoter abolishes erythroid gene expression in Duffy-negative individuals. Nat. Genet. 10, 224–228.

Tsai, A., Muthusamy, A.K., Alves, M.R., Lavis, L.D., Singer, R.H., Stern, D.L., and Crocker, J. (2017). Nuclear microenvironments modulate transcription from low-affinity enhancers. ELife 6, e28975. https://doi.org/10.7554/eLife.28975.

Tuupanen, S., Turunen, M., Lehtonen, R., Hallikas, O., Vanharanta, S., Kivioja, T., Björklund, M., Wei, G., Yan, J., Niittymäki, I., et al. (2009). The common colorectal cancer predisposition SNP rs6983267 at chromosome 8q24 confers potential to enhanced Wnt signaling. Nat. Genet. 41, 885–890. https://doi.org/10.1038/ng.406.

Veerman, C.C., Wilde, A.A.M., and Lodder, E.M. (2015). The cardiac sodium channel gene SCN5A and its gene product NaV1.5: role in physiology and pathophysiology. Gene 573, 177–187. https://doi.org/10.1016/j.gene.2015.08.062.

Visel, A., Rubin, E.M., and Pennacchio, L.A. (2009). Genomic views of distant-acting enhancers. Nature 461, 199–205. https://doi.org/10.1038/nature08451.

Wang, W., Niu, X., Stuart, T., Jullian, E., Mauck, W.M., Kelly, R.G., Satija, R., and Christiaen, L. (2019). A single-cell transcriptional roadmap for cardiopharyngeal fate diversification. Nat. Cell Biol. 21, 674–686. https://doi.org/10.1038/s41556-019-0336-z.

Wang, X., Tucker, N.R., Rizki, G., Mills, R., Krijger, P.H., de Wit, E., Subramanian, V., Bartell, E., Nguyen, X.-X., Ye, J., et al. (2016). Discovery and validation of sub-threshold genome-wide association study loci using epigenomic signatures. ELife 5, e10557. https://doi.org/10.7554/eLife.10557.

Wei, G.-H., Badis, G., Berger, M.F., Kivioja, T., Palin, K., Enge, M., Bonke, M., Jolma, A., Varjosalo, M., Gehrke, A.R., et al. (2010). Genome-wide analysis of ETS-family DNA-binding in vitro and in vivo. EMBO J. 29, 2147–2160. https://doi.org/10.1038/emboj.2010.106.

Woznica, A., Haeussler, M., Starobinska, E., Jemmett, J., Li, Y., Mount, D., and Davidson, B. (2012). Initial deployment of the cardiogenic gene regulatory network in the basal chordate, Ciona intestinalis. Dev. Biol. 368, 127–139. https://doi.org/10.1016/j.ydbio.2012.05.002.

Xie, Y., Su, N., Yang, J., Tan, Q., Huang, S., Jin, M., Ni, Z., Zhang, B., Zhang, D., Luo, F., et al. (2020). FGF/FGFR signaling in health and disease. Signal Transduct. Target. Ther. 5, 181. https://doi.org/10.1038/s41392-020-00222-7.

Ye, M., Coldren, C., Liang, X., Mattina, T., Goldmuntz, E., Benson, D.W., Ivy, D., Perryman, M.B., Garrett-Sinha, L.A., and Grossfeld, P. (2010). Deletion of ETS-1, a gene in the Jacobsen syndrome critical region, causes ventricular septal defects and abnormal ventricular morphology in mice. Hum. Mol. Genet. 19, 648–656. https://doi.org/10.1093/hmg/ddp532.

Zaffran, S., and Frasch, M. (2002). Early Signals in Cardiac Development. Circ. Res. 91, 457–469. https://doi.org/10.1161/01.RES.0000034152.74523.A8.

Zhang, T., Yong, S.L., Tian, X.-L., and Wang, Q.K. (2007). Cardiac-Specific Overexpression of SCN5A Gene Leads to Shorter P Wave Duration and PR Interval in Transgenic Mice. Biochem. Biophys. Res. Commun. 355, 444–450. https://doi.org/10.1016/j.bbrc.2007.01.170.

Zhang, Y., Liu, T., Meyer, C.A., Eeckhoute, J., Johnson, D.S., Bernstein, B.E., Nussbaum, C., Myers, R.M., Brown, M., Li, W., et al. (2008). Model-based Analysis of ChIP-Seq (MACS). Genome Biol. 9, R137. https://doi.org/10.1186/gb-2008-9-9-r137.

